# Mitochondrial RNA capping: highly efficient 5’-RNA capping with NAD^+^ and NADH by yeast and human mitochondrial RNA polymerase

**DOI:** 10.1101/381160

**Authors:** Jeremy G. Bird, Urmimala Basu, David Kuster, Aparna Ramachandran, Ewa Grudzien-Nogalska, Megerditch Kiledjian, Dmitry Temiakov, Smita S. Patel, Richard H. Ebright, Bryce E. Nickels

**Affiliations:** Department of Genetics and Waksman Institute, Rutgers University, USA.; Department of Chemistry and Waksman Institute, Rutgers University, USA.; Department of Biochemistry and Molecular Biology, Rutgers University, Robert Wood Johnson Medical School, USA.; Biochemistry Ph.D. Program, School of Graduate Studies, Rutgers University USA.; Biochemistry Center Heidelberg (BZH), Heidelberg University, Germany.; Department of Cell Biology and Neuroscience, Rutgers University, USA.; Department of Cell Biology, Rowan University, USA.

**Keywords:** mitochondria, RNA capping, NAD^+^, NADH, metabolism, metabolism-transcription coupling, RNA polymerase, transcription, transcription initiation, non-canonical initiating nucleotide, *Saccharomyces cerevisiae*, human cells

## Abstract

Bacterial and eukaryotic nuclear RNA polymerases (RNAPs) cap RNA with the oxidized and reduced forms of the metabolic effector nicotinamide adenine dinucleotide, NAD^+^ and NADH, using NAD^+^ and NADH as non-canonical initiating nucleotides for transcription initiation. Here, we show that mitochondrial RNAPs (mtRNAPs) cap RNA with NAD^+^ and NADH, and do so more efficiently than nuclear RNAPs. Direct quantitation of NAD^+^- and NADH-capped RNA demonstrates remarkably high levels of capping *in vivo*: up to ~60% NAD^+^ and NADH capping of yeast mitochondrial transcripts, and up to ~10% NAD^+^ capping of human mitochondrial transcripts. The capping efficiency is determined by promoter sequence at, and upstream of, the transcription start site and, in yeast and human cells, by intracellular NAD^+^ and NADH levels. Our findings indicate mtRNAPs serve as both sensors and actuators in coupling cellular metabolism to mitochondrial gene expression, sensing NAD^+^ and NADH levels and adjusting transcriptional outputs accordingly.

## Introduction

Chemical modifications of the RNA 5’-end provide a layer of “epitranscriptomic” regulation, influencing RNA fate, including stability, processing, localization, and translation efficiency (1-4). One well-characterized RNA 5’-end modification is the “cap” comprising 7-methylguanylate (m^7^G) added to many eukaryotic messenger RNAs (5-8). Recently, a new RNA 5’-end cap comprising the metabolic effector nicotinamide adenine dinucleotide (NAD) has been shown to be added to certain RNAs isolated from bacterial, yeast, and human cells (9-13).

In contrast to a m^7^G cap, which is added to nascent RNA by a capping complex that associates with eukaryotic RNA polymerase II (RNAP II) (2, 6, 14-16), an NAD cap is added by RNAP itself during transcription initiation, by serving as a non-canonical initiating nucleotide (NCIN) (17) [reviewed in (18, 19)]. NCIN-mediated NAD capping has been demonstrated for bacterial RNAP (13, 17, 20, 21) and eukaryotic RNAP II (17). Thus, whereas m^7^G capping occurs after transcription initiation, on formation of the ~20th RNA bond, and occurs only in organisms harboring specialized capping complexes, NAD capping occurs in transcription initiation, on formation of the first RNA bond, and because it is performed by RNAP itself, is likely to occur in most, if not all, organisms.

NAD exists in oxidized and reduced forms: NAD^+^ and NADH, respectively (Figure 1A). Capping with NAD^+^ has been demonstrated both *in vitro* and *in vivo* (13, 17, 20, 21). Capping with NADH has been demonstrated *in vitro* (17, 20).

**Figure 1.**
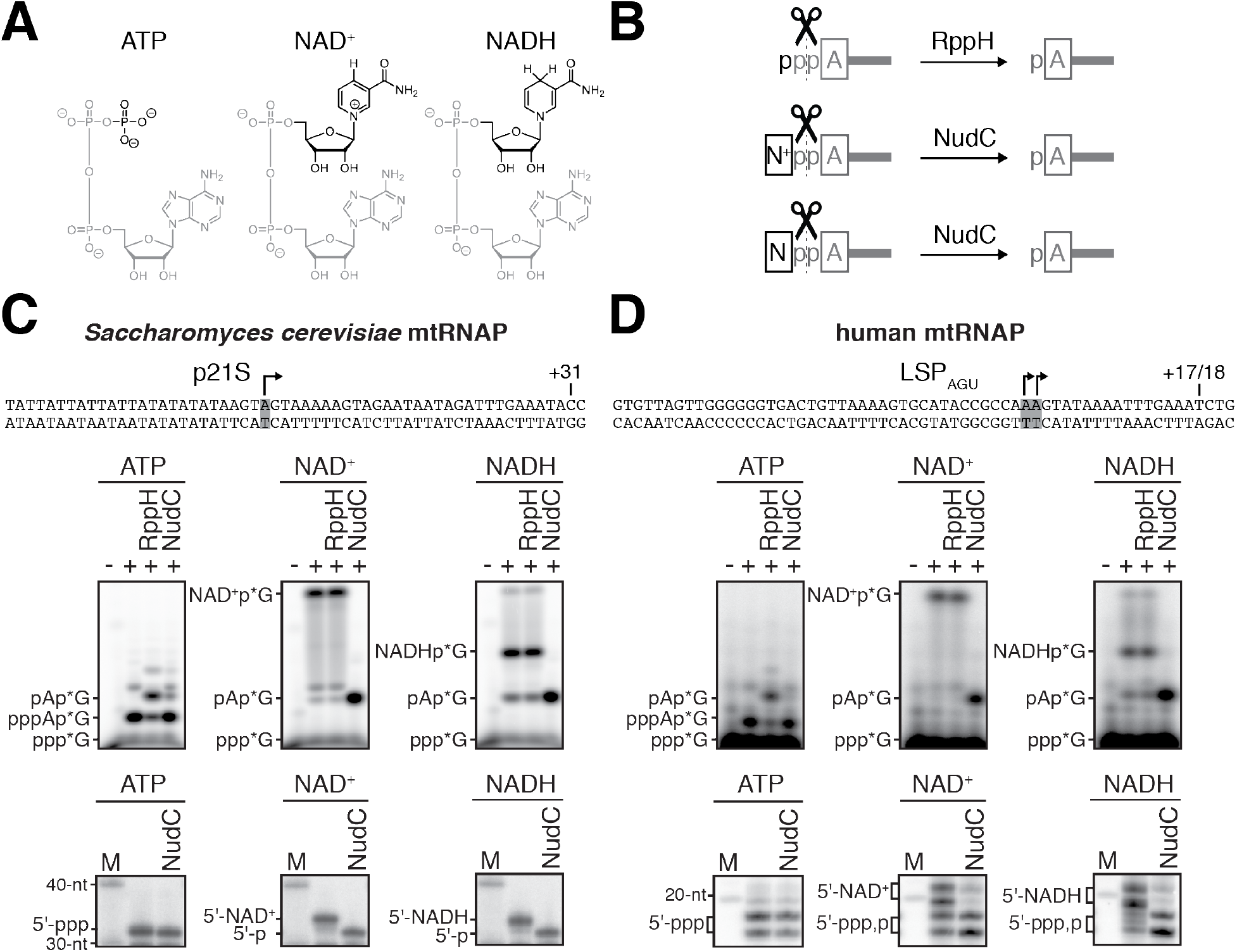
*S. cerevisiae* and human mtRNAPs cap RNA with NAD^+^ and NADH *in vitro*. **A.** Structures of ATP, NAD^+^, and NADH. Grey, identical atoms; black, distinct atoms. **B.** Processing of RNA 5’-ends by RppH and NudC. A, adenosine; N^+^, NAD^+^ nicotinamide; N, NADH nicotinamide; p, phosphate. **C. and D.** NCIN capping with NAD^+^ and NADH by *S. cerevisiae* mtRNAP (C) and human mtRNAP (D). Top, promoter derivatives. Middle, initial RNA products of *in vitro* transcription reactions with ATP, NAD^+^, or NADH as initiating nucleotide and [α^32^P]-GTP as extending nucleotide. Bottom, full-length RNA products of *in vitro* transcription reactions with ATP, NAD^+^, or NADH as initiating nucleotide and [α^32^P]-GTP, ATP, UTP, and 3’-deoxy-CTP (C), or [α^32^P]-GTP, ATP, and UTP (D) as extending nucleotides. Products were treated with RppH or NudC as indicated. Grey box and arrow, transcription start site (TSS); +31 and +17/18, position of last NTP incorporated into RNA; M, 10-nt marker.

Jäschke and co-workers developed a method that combines click-chemistry-mediated covalent capture and high-throughput sequencing, “NAD captureSeq,” to detect NAD^+^-capped RNA (12, 22). Jäschke and co-workers used this method to identify NAD^+^-capped RNAs in bacterial cells [*Escherichia coli* and *Bacillus subtilis*; (12, 13)]. Parker, Kiledjian, and co-workers used the same method to identify NAD^+^-capped RNAs in eukaryotic cells [*Saccharomyces cerevisiae* and human cell line HEK293T; (10, 11)]. Notably, the identified *Saccharomyces cerevisiae* NAD^+^-capped RNAs included not only RNAs produced by nuclear RNAPs, but also RNAs produced by mitochondrial RNAP (mtRNAP). The eukaryotic nuclear RNAPs–RNAP I, II, and III–are multi-subunit RNAPs closely related in sequence and structure to bacterial RNAP (23-26); in contrast, mtRNAPs are single-subunit RNAPs that are unrelated in sequence and structure to multi-subunit RNAPs and, instead, are related to DNA polymerases, reverse transcriptases, and DNA-dependent RNAPs from T7-like bacteriophages (27-33).

The identification of NAD^+^-capped mitochondrial RNAs in *S. cerevisiae* raises the question of whether eukaryotic single-subunit mtRNAPs–like the structurally unrelated bacterial and eukaryotic nuclear multi-subunit RNAPs–can perform NCIN-mediated capping. A recent review discussed evidence supporting the hypothesis that human mtRNAP can perform NCIN capping (18). Here, we show that single-subunit *S. cerevisiae* mtRNAP and human mtRNAP perform NCIN-mediated capping with NAD^+^ and NADH *in vitro*, and do so substantially more efficiently than bacterial and eukaryotic multi-subunit RNAPs. Further, we show that capping efficiency is determined by promoter sequence, we demonstrate very high levels of NAD^+^ and NADH capping–up to ~50%–of mitochondrial transcripts *in vivo*, and we demonstrate that the extents of capping *in vivo*, and distributions of NAD^+^ capping vs. NADH capping *in vivo* are influenced by intracellular levels of NAD^+^ and NADH.

## Results

### S. cerevisiae and human mtRNAPs cap RNA with NAD^+^ and NADH in vitro

To assess whether mtRNAP can cap RNA with NAD^+^ and NADH, we performed *in vitro* transcription experiments (Figures 1 and S1). We analyzed *S. cerevisiae* mtRNAP and a DNA template carrying the *S. cerevisiae* mitochondrial 21S promoter (34), and, in parallel, human mtRNAP and a DNA template containing a derivative of the human mitochondrial light-strand promoter, LSP_AGU_ (35) (Figure 1C-D, top). We performed reactions using either ATP, NAD^+^, or NADH as the initiating entity and using [α^32^P]-GTP as the extending nucleotide (Figure 1C-D, middle). We observed efficient formation of an initial RNA product in all cases (Figure 1C-D, middle). The initial RNA products obtained with ATP, but not with NAD^+^ or NADH, were processed by RppH, which previous work has shown to process 5’-triphosphate RNAs to 5’-monophosphate RNAs (36) (Figure 1B), whereas the initial RNA products obtained with NAD^+^ or NADH, but not with ATP, were processed by NudC, which previous work has shown to process 5’-NAD^+^- and 5’-NADH-capped RNAs to 5’-monophosphate RNAs (12, 37) (Figure 1B). The results establish that *S. cerevisiae* mtRNAP and human mtRNAP are able to generate initial RNA products using NAD^+^ and NADH as NCINs.

We next assessed whether the initial RNA products formed using NAD^+^ and NADH as NCINs can be extended to yield full-length RNA products (Figure 1C-D, bottom). We performed parallel transcription experiments using either ATP, NAD^+^, or NADH as the initiating entity and using [α^32^P]-GTP, ATP, UTP, and 3’-deoxy-CTP (Figure 1C, bottom) or [α^32^P]-GTP, ATP, and UTP (Figure 1D, bottom) as extending nucleotides. We observed efficient formation of full-length RNA products in all cases, and we observed that full-length RNA products obtained with NAD^+^ or NADH, but not with ATP, were sensitive to NudC treatment (Figure 1C-D, bottom). Similar results were obtained in transcription experiments using [α^32^P]-ATP or [^32^P]-NAD^+^ as the initiating entity and using non-radiolabeled extending nucleotides (Figure S1). The results establish that *S. cerevisiae* mtRNAP and human mtRNAP not only generate initial RNA products, but also generate full-length RNA products, using NAD^+^ and NADH as NCINs.

### S. cerevisiae and human mtRNAPs cap RNA with NAD^+^ and NADH more efficiently than bacterial and nuclear RNAPs

We next determined the relative efficiencies of NCIN-mediated initiation vs. ATP-mediated initiation, (k_cat_/K_M_)_NCIN_ / (k_cat_/K_M_)_ATP_, for mtRNAPs (Figures 2 and S2). We performed reactions with *S. cerevisiae* mtRNAP and DNA templates carrying the *S. cerevisiae* mitochondrial 21S promoter or 15S promoter (Figures 2A and S2A), and, in parallel, with human mtRNAP and DNA templates carrying the human mitochondrial light-strand promoter (LSP) or heavy-strand promoter (HSP1) (Figures 2B and S2B). We obtained values of (k_cat_/K_M_)_NCIN_ / (k_cat_/K_M_)_ATP_ of ~0.3 to ~0.4 for NCIN-mediated initiation with NAD^+^ and NADH by *S. cerevisiae* mtRNAP and ~0.2 to ~0.6 for NCIN-mediated initiation with NAD^+^ and NADH by human mtRNAP. These values imply that NCIN-mediated initiation with NAD^+^ or NADH is up to 40% as efficient as initiation with ATP for *S. cerevisiae* mtRNAP and up to 60% as efficient as initiation with ATP for human mtRNAP.

**Figure 2.**
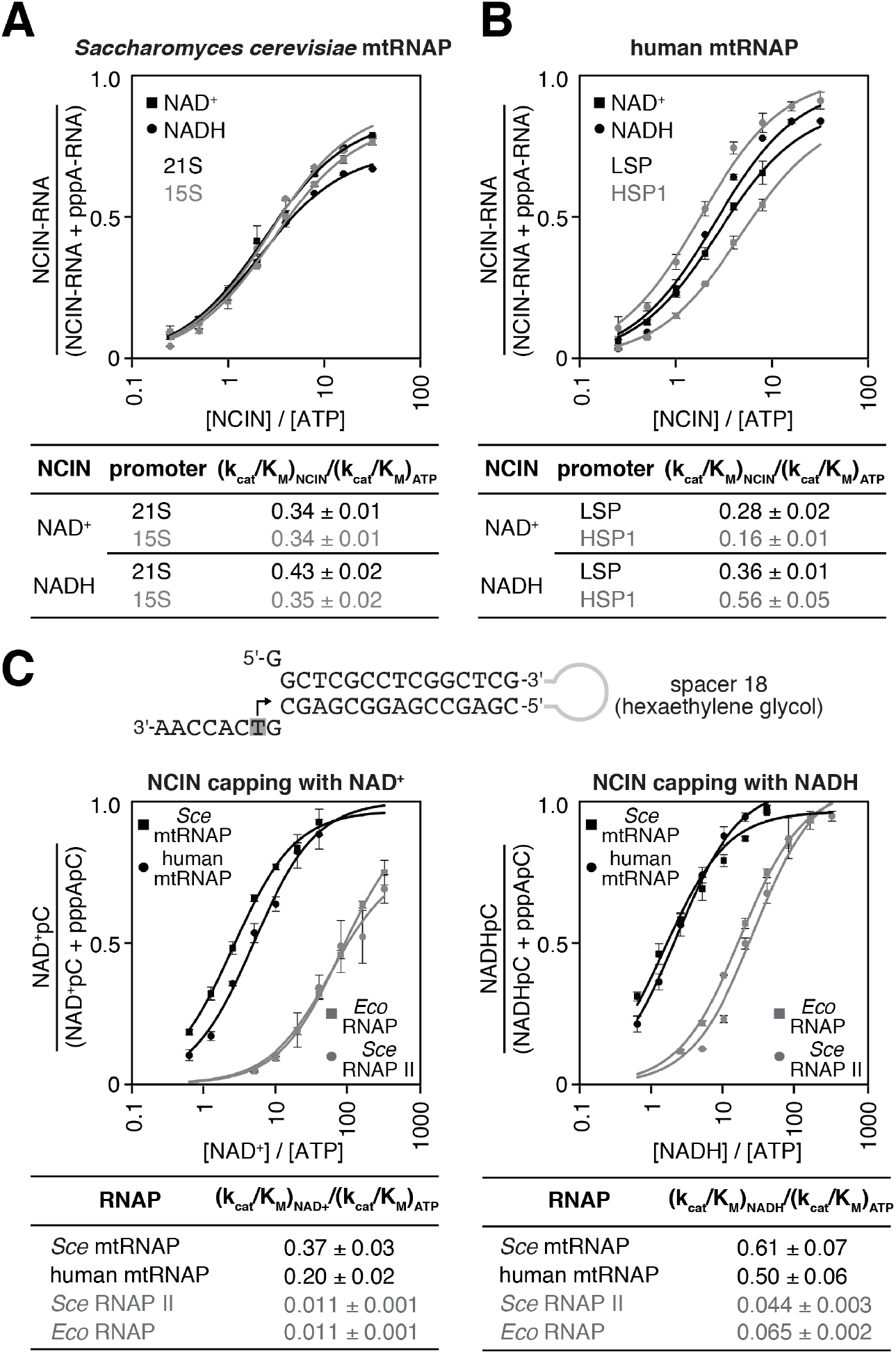
*S. cerevisiae* and human mtRNAPs cap RNA with NAD^+^ and NADH more efficiently than bacterial and nuclear RNAPs. **A. and B.** Dependence of NCIN-mediated capping with NAD^+^ and NADH on [NCIN] / [ATP] ratio for *S. cerevisiae* mtRNAP (A) and human mtRNAP (B) (mean±SD; n=3). DNA templates and representative data are shown in Figure S2. **C.** Dependence of NCIN-mediated capping with NAD^+^ and NADH on [NCIN] / [ATP] ratio for mtRNAPs vs. nuclear RNAPs. Top, Fork junction template. Grey box and arrow indicate TSS. Bottom, Dependence of NCIN-mediated capping with NAD^+^ and NADH on [NCIN] / [ATP] ratio for *S. cerevisiae* mtRNAP (*Sce* mtRNAP), human mtRNAP, *E. coli* RNAP (*Eco* RNAP) and *S. cerevisiae* RNAP II (*Sce* RNAP II) (mean±SD; n=3).

The observed efficiencies of NCIN-mediated initiation with NAD^+^ or NADH by mtRNAPs are substantially higher than the highest previously reported efficiencies for NCIN-mediated initiation with NAD^+^ or NADH by cellular RNAPs [~15%; (17, 21, 38)]. To enable direct comparison of efficiencies of NCIN capping by mtRNAPs vs. cellular RNAPs on the same templates under identical reaction conditions, we performed transcription assays using a “fork-junction” template that bypasses the requirement for sequence-specific RNAP-DNA interactions and transcription-initiation factor-DNA interactions for transcription initiation (Figure 2C, top). In these experiments, we observe efficiencies of NCIN-mediated initiation with NAD^+^ and NADH by mtRNAP that are fully ~10- to ~40-fold higher than efficiencies of NCIN-mediated initiation with NAD^+^ and NADH by *E. coli* RNAP and *S. cerevisiae* RNAP II (Figure 2C, bottom). We conclude that *S. cerevisiae* mtRNAP and human mtRNAP cap RNA with NAD^+^ and NADH more efficiently than bacterial RNAP and eukaryotic nuclear RNAP II.

We next used the same fork-junction template and reaction conditions as in assays performed with mtRNAPs to determine the efficiency of NCIN-mediated initiation with NAD^+^ and NADH for the single-subunit RNAP of bacteriophage T7 (T7 RNAP) (Figure S3). The efficiencies of NCIN-mediated initiation with NAD^+^ and NADH by T7 RNAP were nearly as high as the efficiencies of NCIN-mediated initiation by mtRNAPs. We conclude that there is a quantitative difference in the efficiency of NCIN capping between members of the single-subunit RNAP family (T7 RNAP and mtRNAPs) and members of the multi-subunit RNAP family (bacterial RNAP and eukaryotic nuclear RNAP II).

### Promoter sequence determines efficiency of RNA capping by mtRNAP

In previous work, we have shown that NCIN capping with NAD^+^ and NADH by bacterial RNAP is determined by promoter sequence, particularly at and immediately upstream of, the transcription start site (TSS) (17, 21). NCIN capping by bacterial RNAP occurs only at promoters where the base pair (nontemplate-strand base:template-strand base) at the TSS is a A:T (+1A promoters), and occurs most efficiently at the subset of +1A promoters where the base pair immediately upstream of the TSS is purine:pyrimidine (-1R promoters). We have further shown that sequence determinants for NCIN capping by bacterial RNAP reside within the template strand of promoter DNA (i.e., the strand that templates incoming nucleotide substrates) (21).

To determine whether the specificity for A:T at the TSS (position +1), observed with bacterial RNAP, also is observed with mtRNAP, we assessed NAD^+^ capping by *S. cerevisiae* mtRNAP using promoter derivatives having A:T or G:C at position +1 (Figure 3A-B). We observed NAD^+^ capping in reactions performed using the promoter derivative having A:T at position +1, but not in reactions performed using the promoter derivative having G:C at position +1 (Figure 3B), indicating specificity for A:T at position +1. To determine whether specificity resides in the template strand for A:T at position +1, we analyzed NAD^+^ capping with *S. cerevisiae* mtRNAP using template derivatives having noncomplementary nontemplate- and template-strand-nucleotides (A/C or G/T) at position +1 (Figure 3B). We observed NAD^+^ capping only with the promoter derivative having T as the template strand base at position +1, indicating that specificity at position +1 resides in the template strand.

**Figure 3.**
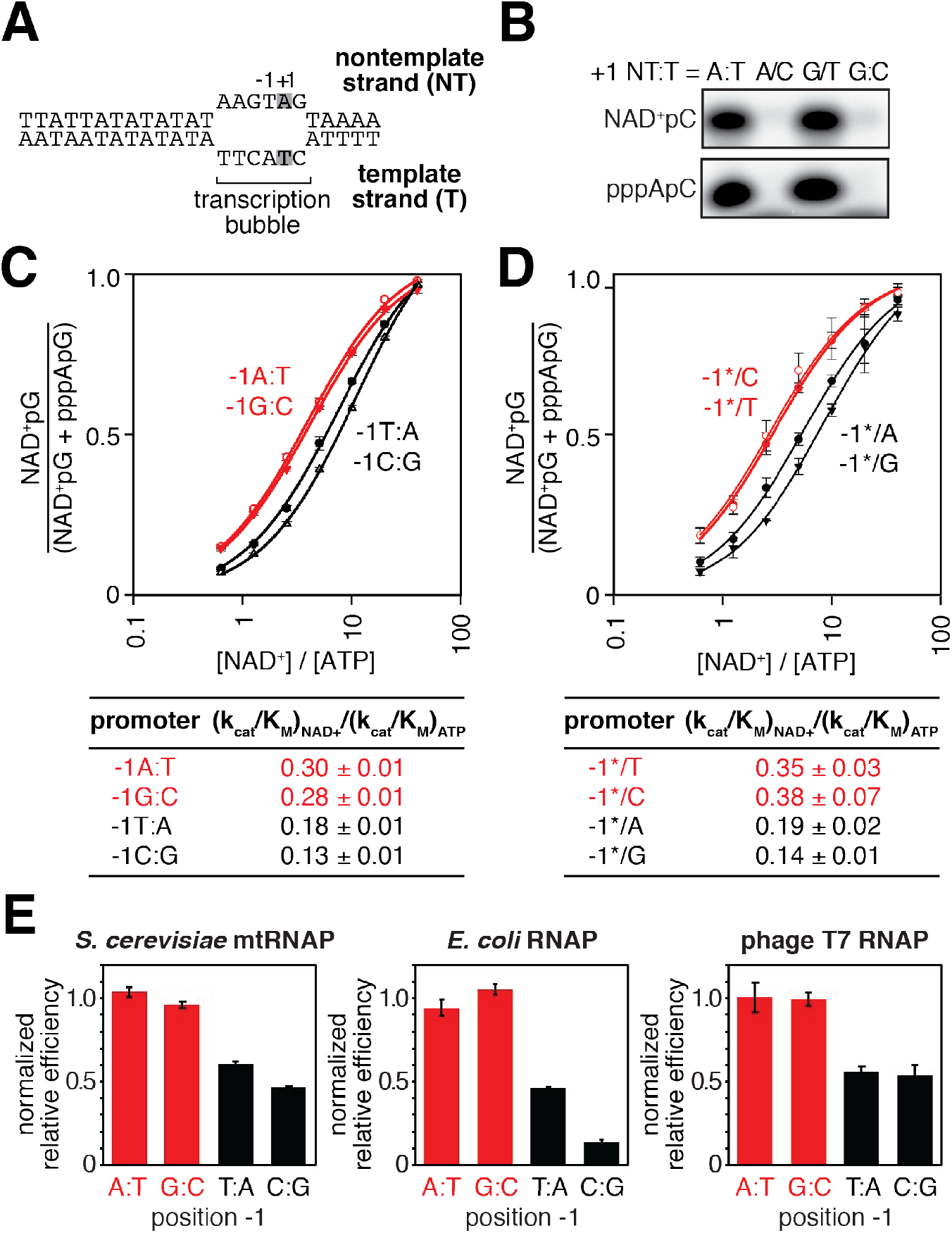
Promoter sequence determines efficiency of RNA capping with NAD^+^ by mtRNAP. **A.** *S. cerevisiae* mitochondrial 21S promoter DNA depicted in the context of the mtRNAP-promoter open complex. DNA non-template strand (NT) on top; DNA template strand (T) on bottom; Unwound, non-base-paired DNA region, “transcription bubble,” indicated by raised and lowered nucleotides; +1 and grey box, bases at the TSS; -1, bases immediately upstream of the TSS. **B.** Products of transcription reactions with NAD^+^ as initiating nucleotide and [α^32^P]-CTP as extending nucleotide for templates having complementary or non-complementary nucleotides at position +1. **C.** Dependence of NAD^+^ capping on [NAD^+^] / [ATP] ratio for homoduplex templates having A:T, G:C, T:A, or C:G at position -1 relative to TSS (mean±SD; n=3). Red, -1R promoters; black, -1Y promoters. **D.** Dependence of NAD^+^ capping on [ATP] / [NAD^+^] ratio for heteroduplex templates having an abasic site (*) on the DNA non-template strand (mean±SD; n=3). Red, promoters with a template-strand Y; black, promoters with a template-strand R. **E.** Sequence preferences at position -1 for *S. cerevisiae* mtRNAP, *E. coli* RNAP, and T7 RNAP. Graphs show normalized values of (kcat/KM)NAD+ / (kcat/KM)ATP determined for homoduplex templates having A:T, G:C, T:A, or C:G at position -1 (mean±SD; n=3). Normalized values were calculated by dividing the value for each individual promoter by the average value measured for -1R promoters. Data for *S. cerevisiae* mtRNAP is from panel C, data for *E. coli* RNAP is from (21), and data for T7 RNAP is from Figure S4.

To determine whether specificity for R:Y at position -1, observed with bacterial RNAP, also is observed with mtRNAP, we analyzed NAD^+^ capping by *S. cerevisiae* mtRNAP using promoter derivatives having either R:Y (A:T or G:C) or Y:R (C:G or T:A) at position -1 (Figure 3C). We observed higher efficiencies of NAD^+^ capping with promoter derivatives having R:Y at position -1 than with promoter derivatives having Y:R (Figure 3C). To determine whether specificity at position -1 resides in the DNA template strand, we performed experiments using promoter derivatives having Y (C or T) or R (A or G) at position -1 of the template strand and having an abasic site (*) on the nontemplate strand (Figure 3D). We observed higher efficiencies of NAD^+^ capping in reactions performed using promoter derivatives having Y at template-strand position -1 than with those having R. Furthermore, within error, the capping efficiencies for promoter derivatives having Y or R at template-strand position -1 matched the capping efficiencies for homoduplex promoter derivatives (Figure 3C-D), indicating that sequence information for NAD^+^ capping with *S. cerevisiae* mtRNAP resides exclusively in the template strand.

We conclude that NCIN capping with NAD^+^ by mtRNAP is determined by the sequence at, and immediately upstream of, the TSS (positions +1 and -1, respectively). We further conclude that the sequence and strand preferences at positions +1 and -1 for NCIN capping with NAD^+^ by mtRNAP match the sequence and strand preferences observed for bacterial RNAP (Figure 3C-E) (17, 21), suggesting that these sequence and strand preferences may be universal determinants of NCIN capping with NAD^+^ for all RNAPs. Consistent with this hypothesis, we find that sequence preferences for NCIN capping with NAD^+^ by bacteriophage T7 RNAP, another member of the single-subunit RNAP family, match the sequence preferences observed for *S. cerevisiae* mtRNAP and bacterial RNAP (Figures 3E and S4). Further consistent with this hypothesis, structural modeling suggests the basis for these sequence and strand preferences is universal: specifically, a requirement for template-strand +1T for base pairing to the NAD^+^ adenine moiety, and a requirement for template strand -1Y for “pseudo” base pairing to the NAD^+^ nicotinamide moiety (17, 21).

### Detection and quantitation of NAD^+^- and NADH-capped mitochondrial RNA in vivo: boronate affinity electrophoresis with processed RNA and synthetic standards

Kössel, Jäschke and co-workers have demonstrated that boronate affinity electrophoresis allows resolution of uncapped RNAs from capped RNAs–such as m^7^G, NAD^+^ and NADH–that contain a vicinal-diol moiety (39-41). However, the procedures of Kössel, Jäschke and co-workers have two limitations: (i) boronate affinity electrophoresis does not allow resolution of RNAs longer than ~200 nt, and (ii) boronate affinity electrophoresis, by itself, is unable to distinguish between different vicinal-diol containing cap structures (m^7^G, NAD^+^, NADH, or others). Here, to overcome these limitations, we have combined boronate affinity electrophoresis with use of oligodeoxynucleotide-mediated RNA cleavage (“DNAzyme” cleavage) (42) and use of synthetic NCIN-capped RNA standards generated using NCIN-mediated transcription initiation *in vitro* (Figure 4). Use of DNAzyme cleavage enables processing of long RNAs to yield defined, short, 5’-end-containing subfragments (Figure 4A). Use of synthetic NCIN-capped RNA standards enables distinction between capped species (Figure 4A,D).

**Figure 4.**
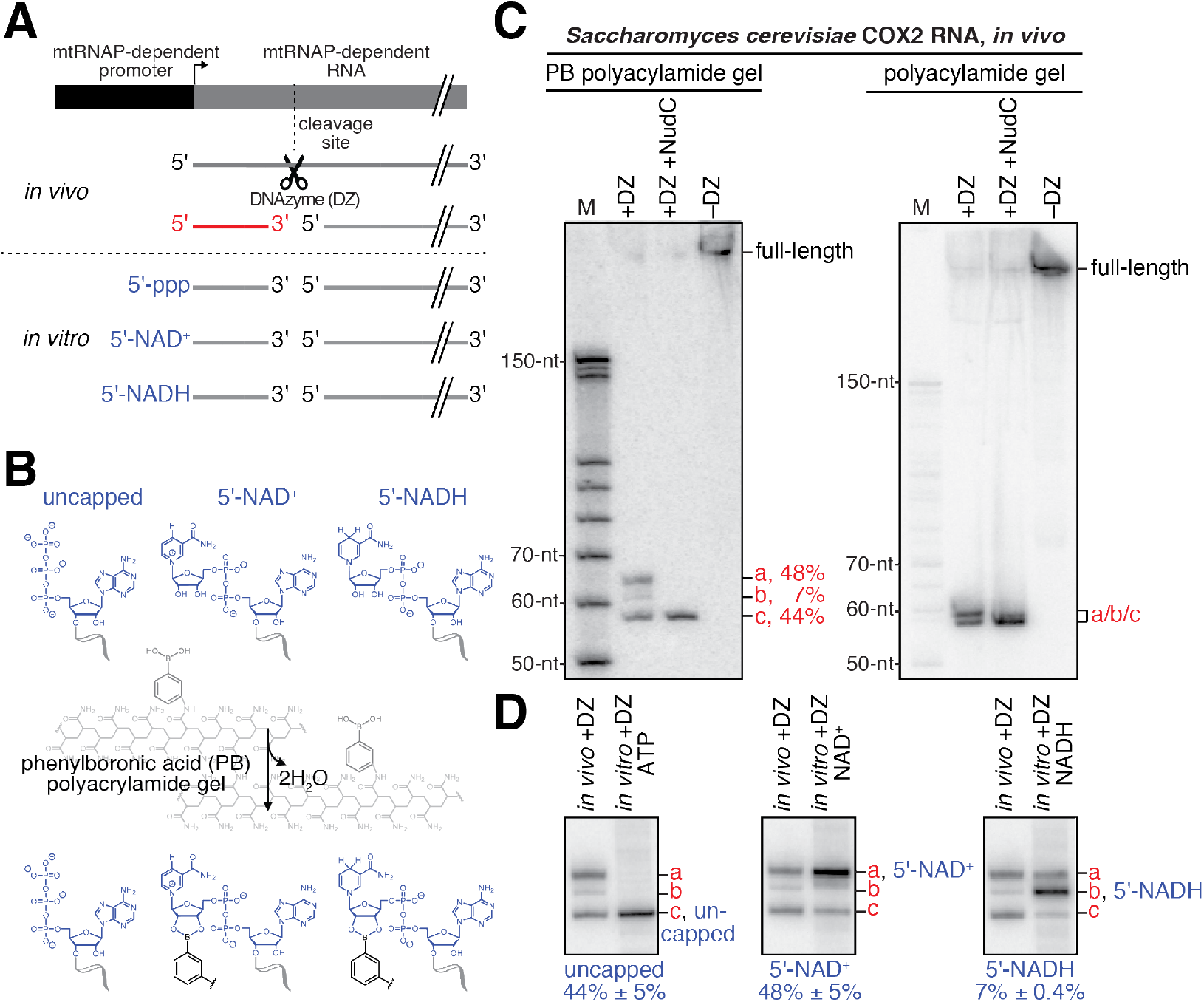
Detection and quantitation of NAD^+^- and NADH-capped mitochondrial RNA *in vivo*: boronate affinity electrophoresis with DNAzyme-cleaved cellular RNA and DNAzyme-cleaved synthetic NCIN-capped RNA standards. **A.** Use of DNAzyme (DZ) to process RNA to yield a defined, short 5’-end-containing subfragment, in parallel *in vivo* (red) and *in vitro* (blue). Uncapped, 5’-triphosphate (ppp) end generated using ATP-mediated initiation; 5’-NAD^+^, NAD^+^-capped end generated using NAD^+^-mediated initiation; 5’-NADH, NADH-capped end generated using NADH -mediated initiation. **B.** Use of boronate affinity electrophoresis to resolve 5’-uncapped, 5’-NAD^+^, and 5’-NADH containing RNAs. Grey, structure of phenylboronic acid (PB) polyacrylamide gel. **C.** PB-polyacylamide gel (left) and polyacrylamide gel (right) analysis of DNAzyme-generated 5’-end-containing subfragments of *S. cerevisiae* mitochondrial RNA COX2. Red, observed 5’-end-containing RNA subfragments resolved by PB-polyacylamide-gel (left) or not resolved by polyacrylamide gel (right); identities of these subfragments are defined in Panel D. **D.** Comparison of electrophoretic mobilities of observed 5’-end-containing subfragments of COX2 RNA generated *in vivo* to 5’-end-containing subfragments of synthetic RNA standards generated *in vitro*. a, NAD^+^-capped RNA; b, NADH-capped RNA; c, uncapped RNA (mean±SD; n=3).

To detect and quantify NCIN capping with NAD^+^ and NADH in mitochondrial RNA isolated from cells, we employed the following steps: (i) DNAzyme cleavage of target RNAs to generate 5’-end-containing subfragments < 80 nt in length (Figure 4A, top); (ii) DNAzyme treatment of synthetic NAD^+^- and NADH-capped RNA standards having sequences identical to RNAs of interest (Figure 4A, bottom); (iii) boronate affinity electrophoresis of DNAzyme-generated subfragments of mitochondrial RNA and DNAzyme-generated subfragments of synthetic NAD^+^- and NADH-capped RNA standards having sequences identical to RNAs of interest (Figure 4B); and (iv) detection of DNAzyme-generated subfragments of mitochondrial RNAs and synthetic RNA standards by hybridization with a radiolabeled oligodeoxyribonucleotide probe (Figure 4C-D).

We selected for analysis two *S. cerevisiae* mitochondrial RNAs that previously had been detected as NAD^+^-capped: COX2 and 21S (10). We isolated *S. cerevisiae* total RNA and analyzed COX2 and 21S RNAs using the procedure described in the preceding paragraph. The results are presented in Figures 4C-D and 5A (top). For both COX2 and 21S RNAs, we detect at least one RNA species with electrophoretic mobility retarded as compared to that of uncapped RNA, indicating the presence of capped RNA. Treatment with the decapping enzyme NudC eliminates these species, confirming the presence of capping. Comparison of the electrophoretic mobilities to those of synthetic NAD^+^- and NADH-capped RNA standards indicates that one of the capped species is NAD^+^-capped RNA, and the other capped species, present under these growth conditions only for COX2 RNA, is NADH-capped RNA. The results show that both COX2 and 21S RNAs are present in NAD^+^-capped forms, and that COX2 RNA also is present in an NADH-capped-form. The results confirm that *S. cerevisiae* mitochondrial RNAs undergo NAD^+^ capping in cells, show that *S. cerevisiae* mitochondrial RNAs undergo NAD^+^ capping at 5’ ends generated by transcription initiation (as opposed 5’ ends generated by RNA processing), and show that *S. cerevisiae* mitochondrial RNAs also undergo NADH capping in cells.

The observed levels of NAD^+^- and NADH-capping of *S. cerevisiae* mitochondrial RNAs are remarkably high. For COX2 RNA, NAD^+^-capped RNA comprises ~50% of the total COX2 RNA pool and NADH-capped RNA comprises ~10% of the total COX2 RNA pool (Figures 4C-D and 5A, top). For 21S RNA, NAD^+^-capped RNA comprises ~30% of the total 21S RNA pool (Figure 5A, top). These levels of NCIN capping are ~2 to ~50 times higher than levels of NCIN-capping in exponentially growing *E. coli* (less than 1% to ~20% for NAD^+^-capping; not previously detected for NADH-capping) (12, 17, 21, 41).

We performed analogous experiments analyzing RNAs produced by transcription from the human mitochondrial LSP promoter (Figure 5B, top). We isolated and analyzed total RNA from HEK293T cells. We observed an NAD^+^-capped species comprising ~10% of the total LSP-derived RNA pool (Figure 5B, top). The results establish that human mitochondrial RNAs undergo NAD^+^ capping in cells and show that human mitochondrial RNAs undergo NAD^+^ capping at 5’ ends generated by transcription initiation (as opposed 5’ ends generated by RNA processing).

**Figure 5.**
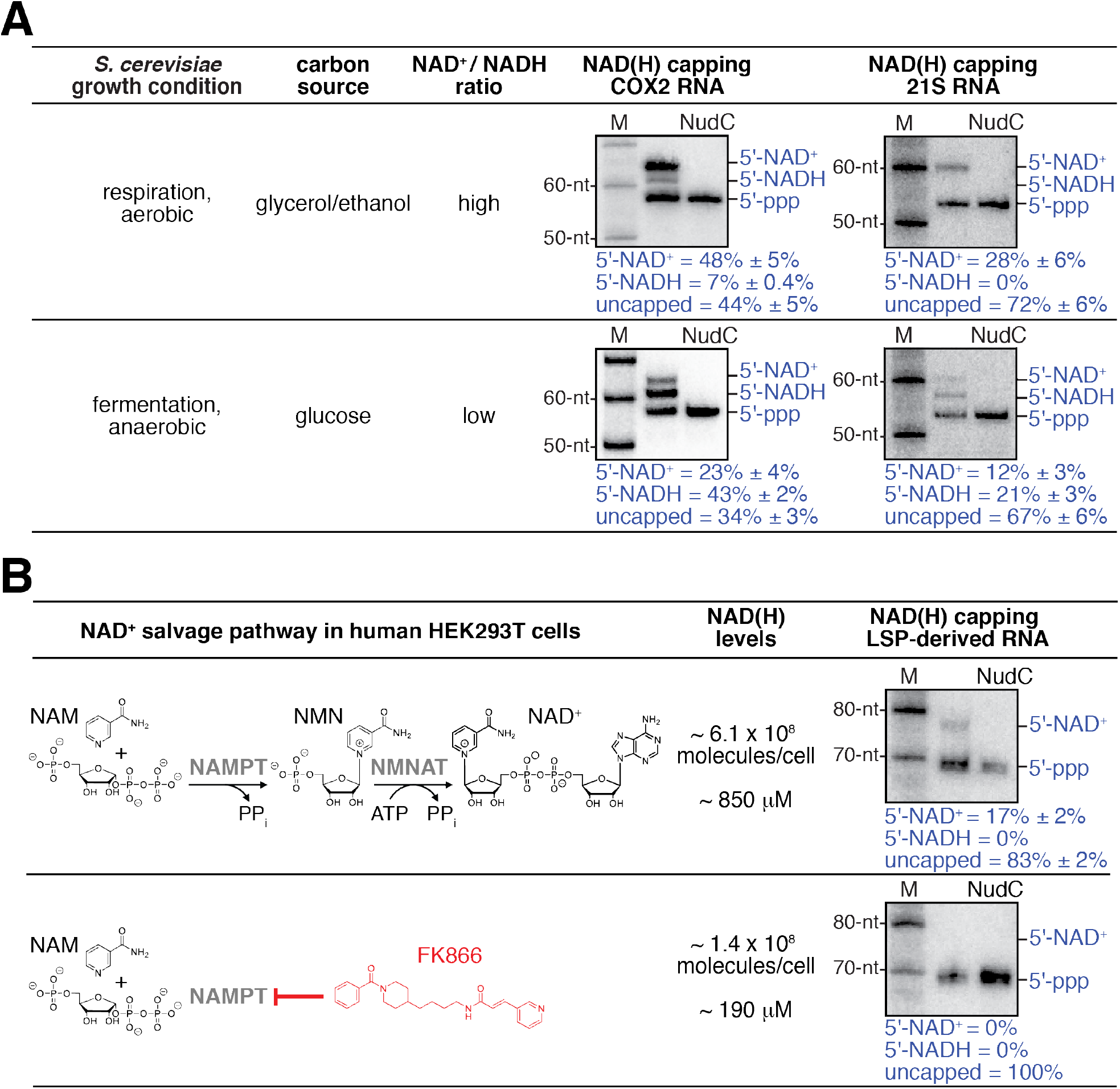
Detection and quantitation of NAD^+^- and NADH-capped mitochondrial RNA *in vivo*: effects of intracellular NAD^+^ and NADH levels in *S. cerevisiae* and human cells. **A.** Changes in intracellular NAD^+^ / NADH ratios result in changes in levels of NAD^+^- and NADH-capped mitochondrial RNA (mean±SD; n=3). Gel images show representative data for *S. cerevisiae* COX2 RNA (left) and 21S RNA (right). Blue annotations as in Figure 4. **B.** Changes in intracellular NAD(H) levels result in changes in levels of NAD^+^- and NADH-capped mitochondrial RNA (mean±SD; n=3). Gel images show representative data for LSP-derived RNAs. Red, NAD(H) biosynthesis inhibitor FK866; NAMPT, Nicotinamide phosphoribosyltransferase; NMNAT, Nicotinamide mononucleotide adenylyltransferase.

### Detection and quantitation of NAD^+^- and NADH-capped mitochondrial RNA in vivo: mtRNAPs serve as both sensors and actuators in coupling cellular metabolism to mitochondrial gene expression

Mitochondria are the primary locus of metabolism and energy transformation in the eukaryotic cell, serving as the venue for the tricarboxylic acid cycle (TCA) cycle and oxidative phosphorylation. The TCA cycle reduces NAD^+^ to NADH and oxidative phosphorylation oxidizes NADH to NAD^+^. Our finding that mtRNAPs perform NCIN capping with NAD^+^ and NADH at efficiencies that vary in a simple mass-action-dependent fashion with [NAD^+^] / [ATP] and [NADH] / [ATP] ratios *in vitro* (Figures 1, 2, S1, and S2), and our finding that mtRNAPs perform efficient NCIN capping to yield NAD^+^- and NADH-capped mitochondrial RNAs *in vivo* (Figures 4 and 5), raise the possibility that mtRNAPs may serve as both sensors and actuators in coupling metabolism to mitochondrial gene expression *in vivo*.

As a first test of this hypothesis, we assessed whether changing intracellular [NAD^+^] / [NADH] ratios results in changes in NAD^+^ and NADH capping of mitochondrial RNAs. We isolated total RNA from *S. cerevisiae* grown under conditions that result either high or low [NAD^+^] / [NADH] ratios (43, 44): i.e., respiration (glycerol/ethanol; aerobic) or fermentation (glucose; anaerobic). We analyzed the same two mitochondrial RNAs as above: COX2 and 21S (Figure 5A). We observed marked changes in levels of NAD^+^ and NADH capping for both analyzed mitochondrial RNAs. For COX2 RNA, on changing from the growth condition yielding a high [NAD^+^] / [NADH] ratio to the growth condition yielding a low [NAD^+^] / [NADH] ratio, we observe a decrease in levels of NAD^+^ capping (from ~50% to ~20%) and an anti-correlated increase in the level of NADH capping (from ~10% to ~40%). Notably, the total level of NAD^+^ and NADH capping, NAD(H) capping, remains constant under the two conditions (~60%), indicating that the relative levels of NCIN-mediated initiation and ATP-mediated initiation do not change (Figure 5A). For 21S RNA, the same pattern is observed (Figure 5A): on changing from the growth condition yielding a high [NAD^+^] / [NADH] ratio to the growth condition yielding a low [NAD^+^] / [NADH] ratio, the level of NAD^+^ capping decreases (from ~30% to ~10%), the level of NADH capping increases (from 0% to ~20%), and the total level of NAD^+^ and NADH capping remains constant (~30%). The results indicate that changing the [NAD^+^] / [NADH] ratio changes transcription outputs *in vivo*.

As a second test of this hypothesis, we assessed whether changing intracellular total NAD(H) levels results in changes in NCIN capping of mitochondrial RNAs (Figure 5B). We isolated RNA from human HEK293T cells grown under conditions yielding either high or low intracellular NAD(H) levels: standard growth media or growth media in the presence of the NAD(H)-biosynthesis inhibitor FK866 (45, 46). We analyzed the same LSP-derived mitochondrial RNA as in the preceding section (Figure 5B). On changing from the growth condition yielding high intracellular NAD(H) levels to the growth condition yielding low NAD(H) levels, we observe a marked change in the total level of NCIN capping (from ~10% to 0%). The results indicate that changing NAD(H) levels changes levels of NCIN-capped mitochondrial RNAs *in vivo*.

Taken together, the results of the two experiments in Figure 5 indicate that mtRNAP serves as both sensor and actuator in coupling [NAD^+^] / [NADH] ratios to relative levels of NAD^+^- and NADH-capped mitochondrial RNAs (Figure 5A), and in coupling total NAD(H) levels to total levels of NCIN-capped mitochondrial RNAs (Figure 5B), thereby coupling cellular metabolism to mitochondrial transcription outputs. We suggest that mtRNAPs serve as sensors through their mass-action-dependence in selecting NAD^+^ vs. NADH vs. ATP as initiating entity during transcription initiation, and serve as actuators by incorporating NAD^+^ vs. NADH vs. ATP at the RNA 5’ end during transcription initiation.

## Discussion

Our results show that *S. cerevisiae* and human mtRNAPs cap RNA with NAD^+^ and NADH (Figures 1 and S1), show that *S. cerevisiae* and human mtRNAPs cap RNA with NAD^+^ and NADH more efficiently than bacterial and eukaryotic nuclear RNAPs (Figures 2 and S2), and show that capping efficiency by mtRNAPs is determined by promoter sequence (Figure 3). Our results further show that the proportions of mitochondrial RNAs that are capped with NAD^+^ and NADH are remarkably high–up to ~50% and up to ~40%, respectively (Figures 4 and 5)–and that these proportions change in response to cellular NAD^+^ and NADH levels (Figure 5).

We and others previously have shown that NCIN capping by cellular RNAPs has functional consequences (11, 12, 17). Our results here showing that *S. cerevisiae* and human mitochondrial RNAs are capped at substantially higher levels than nonmitochondrial RNAs–up to ~50% for analyzed *S. cerevisiae* mitochondrial RNAs and up to ~10% for analyzed human mitochondrial RNAs (Figures 4 and 5)–suggest that NCIN capping in mitochondria occurs at a higher efficiency, and has a higher importance, than NCIN capping in other cellular compartments. Four other considerations support this hypothesis. First, mtRNAPs are substantially more efficient at NAD^+^ and NADH capping than bacterial and eukaryotic nuclear RNAPs (Figure 2C). Second, levels of NAD^+^ and NADH in mitochondria are substantially higher than levels in other cellular compartments (47, 48). Third, all *S. cerevisiae* and human mitochondrial promoters are +1A promoters (18 promoters in *S. cerevisiae* mitochondria; 2 promoters in human mitochondria) (49-51), in contrast to bacterial and eukaryotic nuclear RNAP promoters, for which approximately half are +1A promoters (52-56). Fourth, we observe capping with both NAD^+^ and NADH for mitochondrial RNAs *in vivo* (Figures 4 and 5), whereas, to date, capping has been observed with only NAD^+^ for non-mitochondrial RNAs *in vivo*, raising the possibility that, in mitochondria, but not in other cellular compartments, NAD^+^ and NADH caps dictate different RNA fates and, correspondingly, different transcription outputs.

Our results showing that levels of NAD^+^ and NADH capping by mtRNAP correlate with changes in intracellular levels of NAD^+^ and NADH (Figure 5) indicate that mtRNAP serves as a sensor, sensing [NAD^+^] / [ATP] and [NADH] / [ATP] ratios, when selecting initiating entities and, simultaneously, serves as an actuator by altering RNA 5’ ends when selecting initiating entities. Because all *S. cerevisiae* and human mitochondrial promoters are +1A promoters, this dual sensor/actuator activity of mtRNAP will occur at, and couple metabolism to gene expression at, all *S. cerevisiae* and human mitochondrial promoters. This dual sensor/actuator activity of mtRNAP obviates the need for a dedicated signal-processing machinery for coupling metabolism to gene expression, instead employing a pan-eukaryotic housekeeping enzyme for signal processing. As such, this dual sensor/actuator activity of mtRNAP provides a remarkably economical, parsimonious mechanism of coupling metabolism to gene expression.

## Acknowledgements

Work was supported by AHA 16PRE30400001 (UB), and NIH grants GM067005 (MK), GM118086 (SSP), GM104231 (DT), GM041376 (RHE), and GM118059 (BEN). We thank Atif Towheed and Douglas Wallace (Children’s Hospital of Philadelphia) for discussions and providing samples for related experiments.

## Author Contributions

Conceptualization, DT, SSP, RHE, BEN

Methodology, JGB, UB, DK, AR, DT, SSP

Formal Analysis, JGB, UB, DK

Investigation, JGB, UB, DK, AR, EGN

Writing – Original Draft, RHE, BEN

Writing – Review & Editing, JGB, UB, DK, AR, MK, DT, SSP

Visualization, JGB, UB, DK, SSP, RHE, BEN

Supervision, SSP, RHE, BEN

Project Administration, SSP, RHE, BEN

Funding Acquisition, UB, MK, DT, SSP, RHE, BEN

---

**Figure S1.**
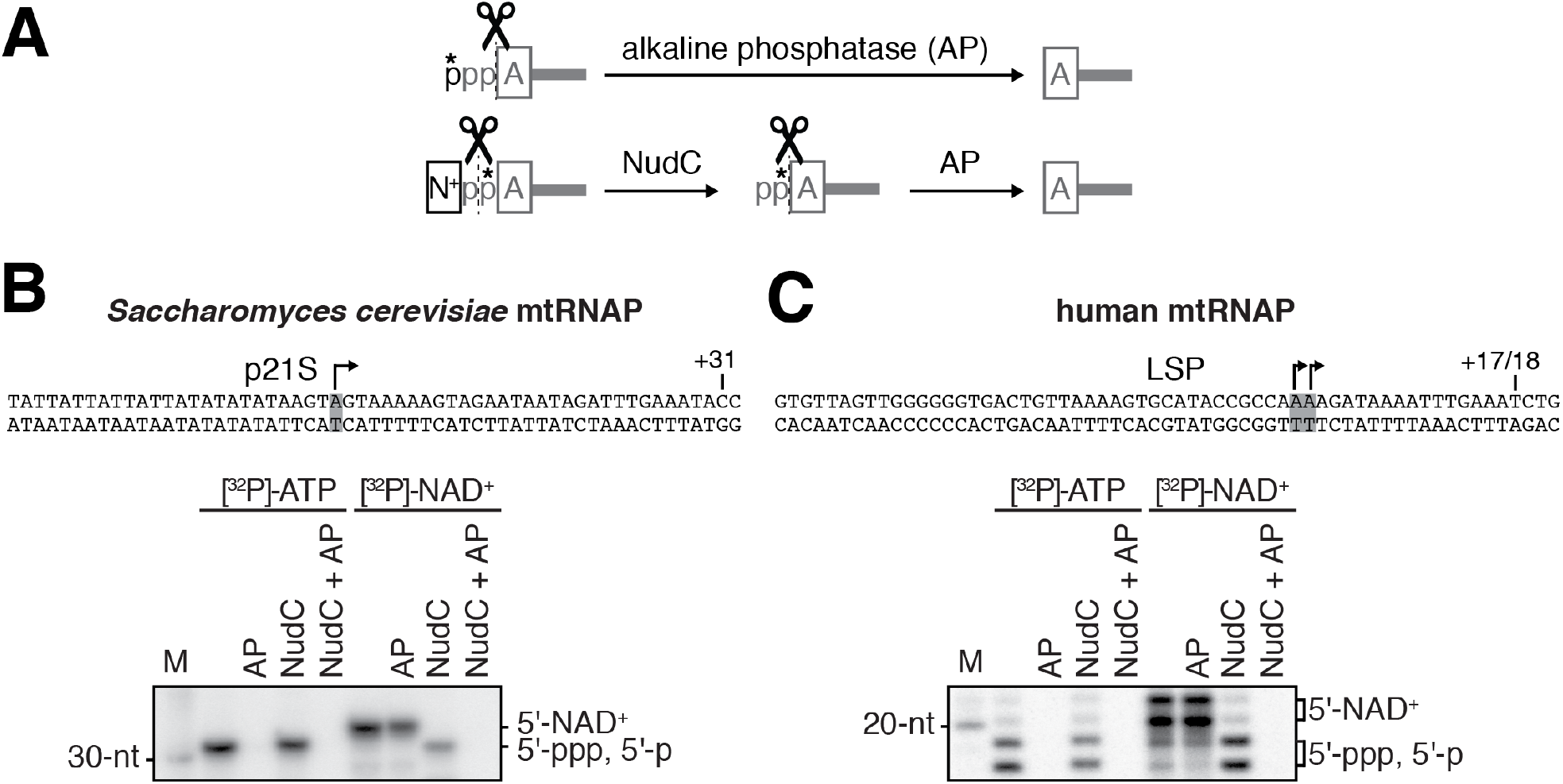
*S. cerevisiae* and human mtRNAPs cap RNA with NAD^+^ *in vitro*: additional data. **A.** Processing of radiolabeled RNA 5’-ends by alkaline phosphatase (AP) and NudC. A, adenosine; N^+^, NAD^+^ nicotinamide; p, phosphate. *, radiolabeled phosphate. **B. and C.** NCIN capping with NAD^+^ by *S. cerevisiae* mtRNAP (B) and human mtRNAP (C). Top, promoter derivatives. Bottom, full-length RNA products of *in vitro* transcription reactions with [γ^32^P]-ATP or [α^32^P]-NAD^+^ as initiating nucleotide and GTP, ATP, UTP, and 3’-deoxy-CTP (C), or GTP, ATP, and UTP (D) as extending nucleotides. Products were treated with NudC alone, AP alone, or NudC and AP, as indicated. Grey box and arrow, TSS; +31 and +17/18, position of last NTP incorporated into RNA; M, 10-nt marker.

**Figure S2.**
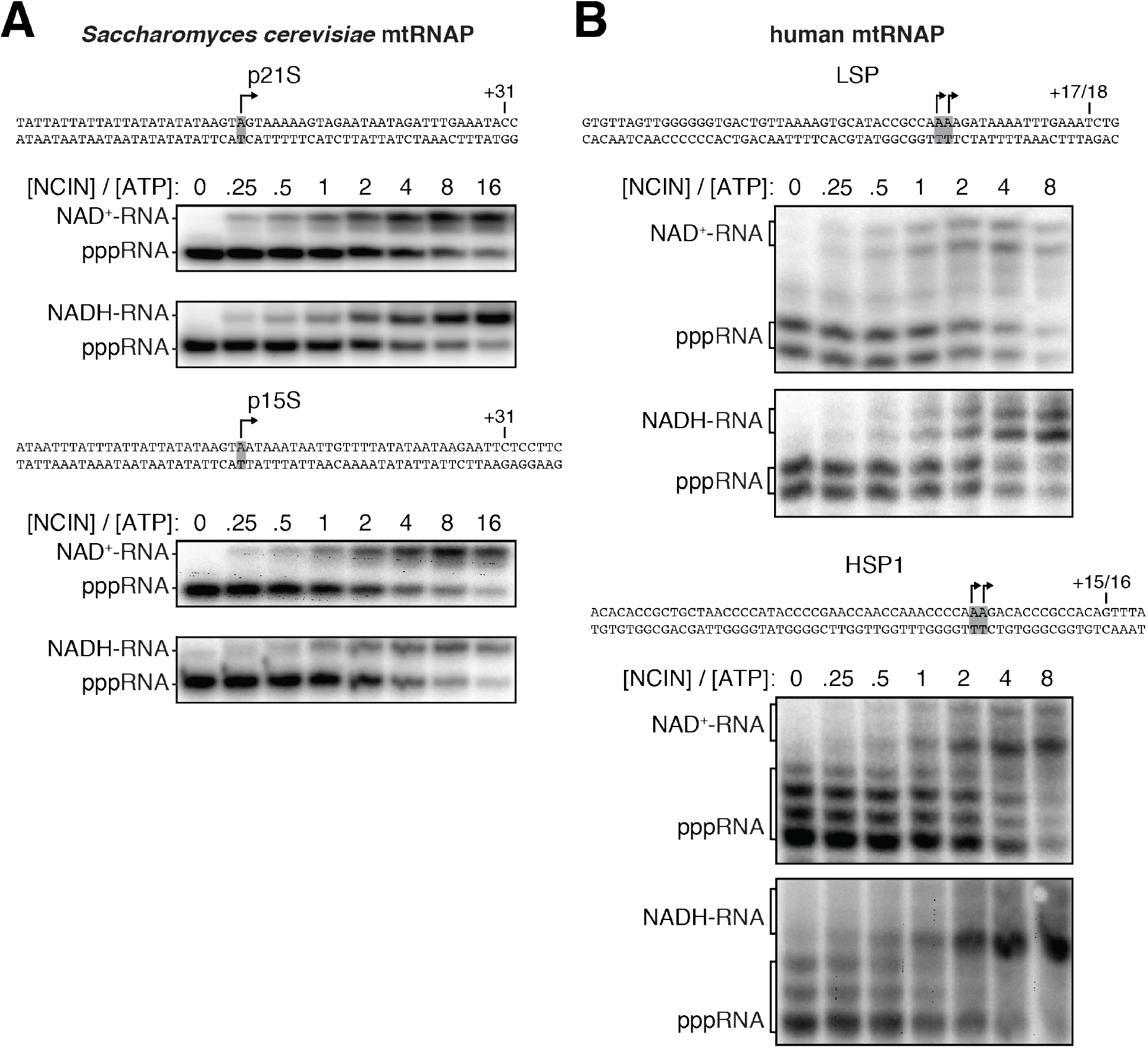
Dependence of NCIN-mediated capping with NAD^+^ and NADH on [NCIN] / [ATP] ratio for mtRNAPs: representative data. **A. and B.** Panels show DNA templates and full-length RNA products of *in vitro* transcription reactions performed with *S. cerevisiae* mtRNAP (A) and human mtRNAP (B) with the indicated [NCIN] / [ATP] ratio. Grey box and arrow, TSS; +31, +17/18, +15/16, position of last NTP incorporated into RNA; M, 10-nt marker.

**Figure S3.**
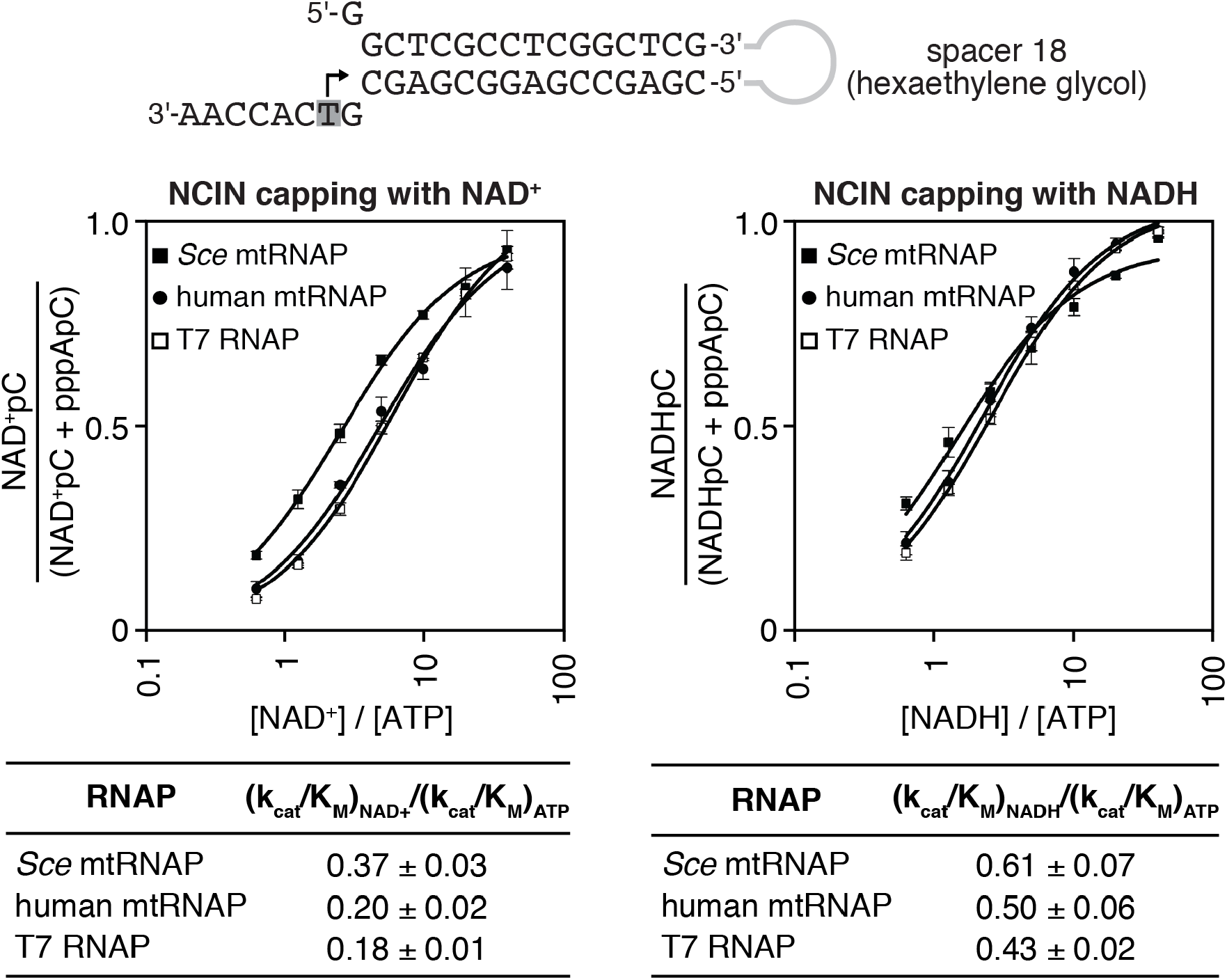
*S. cerevisiae* and human mtRNAPs cap RNA with NAD^+^ and NADH at least as efficiently as bacteriophage T7 RNAP. Dependence of NCIN-mediated capping with NAD^+^ and NADH on [NCIN] / [ATP] ratio for mtRNAPs vs. nuclear RNAPs. Top, Fork junction template. Grey box and arrow indicate TSS. Bottom, Dependence of NCIN-mediated capping with NAD^+^ and NADH on [NCIN] / [ATP] ratio for *S. cerevisiae* mtRNAP (*Sce* mtRNAP), human mtRNAP, and T7 RNAP (mean±SD; n=3).

**Figure S4.**
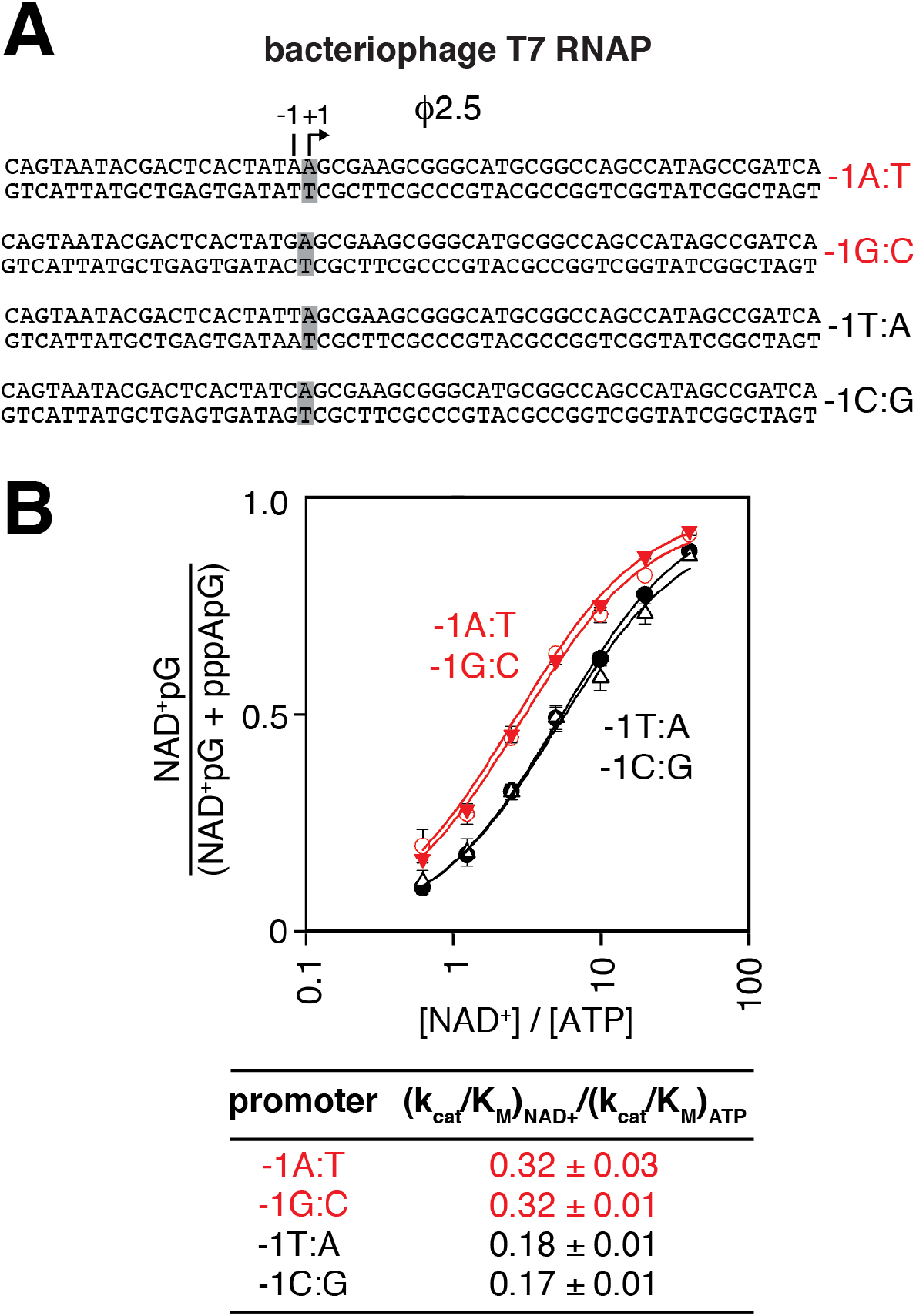
Promoter sequence determines efficiency of RNA capping with NAD^+^: bacteriophage T7 RNAP. **A.** Bacteriophage T7 RNAP-dependent promoter derivatives analyzed. Red, -1R promoters; black, -1Y promoters; +1 and grey box, bases at the TSS; -1, bases immediately upstream of the TSS. **B.** Dependence of NAD^+^ capping on [NAD^+^] / [ATP] ratio for homoduplex templates having A:T, G:C, T:A, or C:G at position -1 relative to TSS (mean±SD; n=3).

**Table S1.**
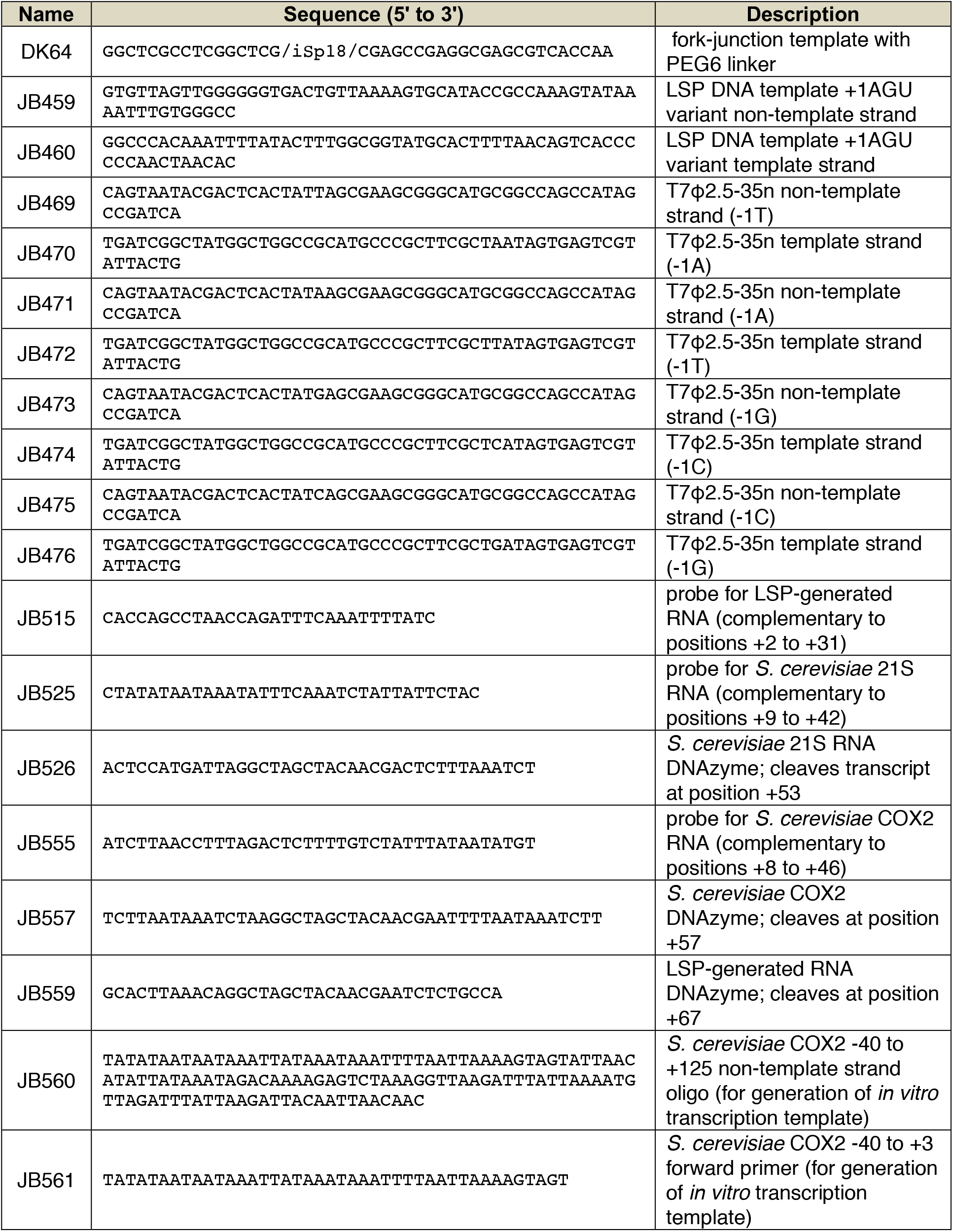
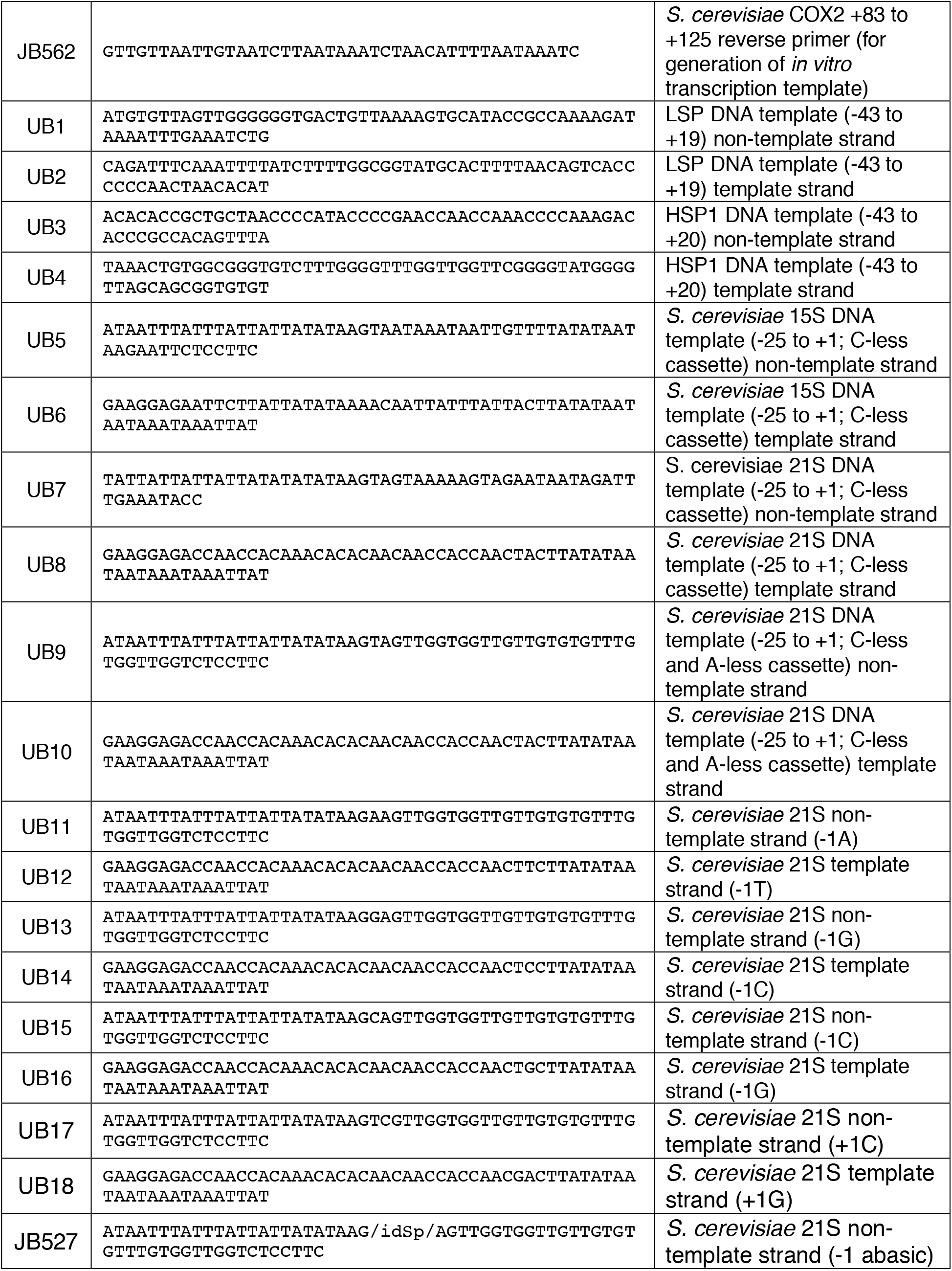
Oligoribodeoxynucleotides.

## Materials and Methods

### Proteins

*S. cerevisiae* mtRNAP (Rpo41) was prepared from *E. coli* strain BL21(DE3) transformed with pJJ1399 (gift of Judith A. Jaehning) using culture and induction procedures, polyethyleneimine treatment, ammonium sulfate precipitation, Ni-sepharose, DEAE sepharose and Heparin-sepharose chromatography as in (57). *S. cerevisiae* Mtf1 was prepared from *E. coli* strain BL21(DE3) transformed with pTrcHisC-Mtf1(58) and purified using culture and induction procedures, polyethyleneimine treatment, ammonium sulfate precipitation, and tandem DEAE sepharose and Ni-sepharose chromatography as in (57).

Human mtRNAP (POLRMT) and TFAM were purified from *E. coli* strain BL21(DE3) transformed with pPROEXHTb-POLRMT(43-1230)-6xHis (59) or pPROEXHTb-TFAM(43-245)-6xHis (59), respectively, using culture and induction procedures, polyethyleneimine treatment, ammonium sulfate precipitation, and Ni-sepharose and heparin-sepharose chromatography as in (59). Human TFB2 was purified from *E. coli* strain ArcticExpress (DE3) (Stratagene) transformed with pT7TEVHMBP4 (60), using culture and induction procedures, polyethyleneimine treatment, ammonium sulfate precipitation, Ni-sepharose and heparin-sepharose chromatography, and size exclusion chromatography as in (60).

T7 RNAP was prepared from *E. coli* strain BL21 transformed with pAR1219 using culture and inductions procedures, SP-Sephadex, CM-Sephadex and DEAE-Sephacel chromatography as described in (61).

*E. coli* RNAP core enzyme was prepared from *E. coli* strain NiCo21(DE3) transformed with plasmid pIA900 (62) using culture and induction procedures, immobilized-metal-ion affinity chromatography on Ni-NTA agarose, and affinity chromatography on Heparin HP as in (62).

*S. cerevisiae* RNA polymerase II core enzyme (gift of Craig Kaplan) was prepared as described in (63).

*E. coli* NudC was prepared from *E. coli* strain NiCo21(DE3) transformed with plasmid pET NudC-His (17) using metal-ion chromatography and size-exclusion chromatography as in (12).

RNA 5’ pyrophosphohydrolase (RppH) and T4 polynucleotide kinase (PNK) were purchased from New England Biolabs (NEB). FastAP Thermosensitive Alkaline Phosphatase was purchased from Thermo Fisher Scientific. Molar concentrations of purified proteins were determined by light absorbance at 280 nm and the calculated respective molar extinction coefficients.

### Oligodeoxyribonucleotides

Sequences of the oligodeoxyribonucleotides used in this work are provided in Table S1. All oligodeoxyribonucleotides were purchased from Integrated DNA Technologies (IDT) with standard desalting purification unless otherwise specified.

Linear *in vitro* transcription templates used for transcription assays shown in Figures 1, 2, 3, S1, S2, S3, and S4 were generated by mixing complementary equimolar amounts of nontemplate- and template-strand DNA in 10 mM Tris HCl pH 8.0, incubating the mixture at 95°C for 5 min, and cooling the mixture by 0.5°C per minute to 25°C.

Transcription templates used to generate *in vitro* RNA standards for Northern analysis (Figures 4 and 5) were generated by PCR. PCR reactions contained a mixture of 5 nM template oligo, 0.5 *µ*M forward primer, 0.5 *µ*M reverse primer, and Phusion HF Master Mix (Thermo Scientific). Reaction products were isolated using a Monarch PCR & DNA cleanup kit (NEB).

The radiolabeled 10-nt marker labeled “M” in gel images shown in Figures 1, 4, 5, and S1 was generated using the Decade Marker System (Thermo Fisher Scientific), PNK (NEB) and [γ^32^P]-ATP (Perkin Elmer; 6,000 Ci/mmol).

### *In vitro* transcription assays

Assays performed with *S. cerevisiae* mtRNAP were based on procedures described in (34). Assays performed with human mtRNAP were based on procedures described in (59).

For initial RNA product assays in Figure 1C, 1 *µ*M DNA template, 1 *µ*M *S. cerevisiae* mtRNAP, and 1 *µ*M Mtf1 were incubated at 25°C for 10 min in *Sce*-mtRNAP reaction buffer (50 mM Tris-acetate pH 7.5, 100 mM potassium glutamate, 10 mM magnesium acetate, 0.01% Tween-20, 1 mM DTT, and 5% glycerol). A mixture containing the initiating nucleotide (200 *µ*M ATP, 1 mM NAD^+^, or 1mM NADH) and extending nucleotide (10 *µ*M of non-radiolabeled GTP plus 6 mCi of [α^32^P]-GTP [Perkin Elmer; 3,000 Ci/mmol]) was added, and assays were incubated at 25°C for 30 min. Radiolabeled initial products were isolated using a Nanosep 3 kDa cutoff spin concentrator (Pall). For Figure 3B, reactions were stopped with 10 *µ*l RNA loading dye (95% deionized formamide, 18 mM EDTA, 0.25% SDS, xylene cyanol, bromophenol blue, amaranth) and were analyzed by electrophoresis on 7.5 M urea, 1x TBE, 20% polyacrylamide gels (UreaGel System; National Diagnostics), followed by storage-phosphor imaging (Typhoon 9400 variable-mode imager; GE Life Science).

For the initial RNA product assays in Figure 1D, 1 *µ*M DNA template, 1 *µ*M human mtRNAP, 1 *µ*M TFAM, and 1 *µ*M TFB2M were incubated at 25°C for 10 min in human-mtRNAP reaction buffer (50 mM Tris-acetate pH 7.5, 50 mM sodium glutamate, 10 mM magnesium acetate, 1 mM DTT, and 0.05% Tween-20). A mixture containing the initiating nucleotide (200 *µ*M ATP, 1 mM NAD^+^, or 1 mM NADH) and extending nucleotide (10 *µ*M of non-radiolabeled GTP plus 6 mCi of [α^32^P]-GTP at [Perkin Elmer; 3,000 Ci/mmol]) was added, and assays were incubated at 25°C for 60 min. Radiolabeled initial RNA products were isolated using a Nanosep 3 kDa cutoff spin concentrator (Pall).

A portion of the recovered initial RNA products were mixed with either 10 U of RppH or 400 nM NudC and incubated at 37°C for 30 min. Reactions were stopped by addition of 10 *µ*l RNA loading dye. Samples were analyzed by electrophoresis on 7.5 M urea, 1x TBE, 20% polyacrylamide gels (UreaGel System; National Diagnostics), followed by storage-phosphor imaging (Typhoon 9400 variable-mode imager; GE Life Science).

For full-length product assays in Figures 1C and S1B, 1 µM DNA template, 1 *µ*M *S. cerevisiae* mtRNAP, and 1 *µ*M Mtf1 were incubated at 25°C for 10 min in *Sce-*mtRNAP reaction buffer. A mixture containing the initiating nucleotide (200 *µ*M ATP, 1 mM NAD^+^, or 1mM NADH for Figure 1C; 200 *µ*M non-radiolabeled ATP plus 10 *µ*Ci [γ^32^P]-ATP [Perkin Elmer; 6,000 Ci/mmol] or 1 mM NAD^+^ plus 20 *µ*Ci [α^32^P]-NAD^+^ [Perkin Elmer; 800 Ci/mmol] for Figure S1B) and extending nucleotides (200 *µ*M GTP, 200 *µ*M 3’-deoxy-CTP, 20 *µ*M ATP, 200 *µ*M non-radiolabeled UTP, and 6 mCi of [α^32^P]-UTP [Perkin Elmer; 3000 Ci/mmol] for Figure 1C; 200 *µ*M GTP, 200 *µ*M 3’-deoxy-CTP, 20 *µ*M ATP, 200 *µ*M UTP for Figure S1B) was added, and assays were incubated at 25°C for 30 min. Reactions were stopped by addition of 1.5x stop solution (0.6 M Tris HCl pH 8.0, 18 mM EDTA, 0.1 mg/ml glycogen), samples were extracted with acid phenol:chloroform (5:1, pH 4.5; Thermo Fisher Scientific), and RNA products were recovered by ethanol precipitation and resuspended in NudC reaction buffer (10 mM Tris HCl pH 8.0, 50 mM NaCl, 10 mM MgCl_2_, 1 mM DTT).

For full-length product assays in Figures 1D and S1C, 1 *µ*M DNA template, 1 *µ*M human mtRNAP, 1 *µ*M TFAM, and 1 *µ*M TFB2M were incubated at 25°C for 10 min in human-mtRNAP reaction buffer. A mixture containing the initiating nucleotide (200 *µ*M ATP, 1 mM NAD^+^, or 1mM NADH for Figure 1D; 200 *µ*M non-radiolabeled ATP plus 10 *µ*Ci [γ^32^P]-ATP [Perkin Elmer; 6000 Ci/mmol] or 1 mM NAD^+^ plus 20 *µ*Ci [α^32^P]-NAD^+^ [Perkin Elmer; 800 Ci/mmol] for Figure S1C) and extending nucleotides (200 *µ*M GTP, 20 *µ*M ATP, 200 *µ*M non-radiolabeled UTP, and 6 mCi of [α^32^P]-UTP [Perkin Elmer; 3000 Ci/mmol] for Figure 1D; 200 *µ*M GTP, 20 *µ*M ATP, 200 *µ*M UTP for Figure S1C) was added, and assays were incubated at 25°C for 60 min. Reactions were stopped by addition of 1.5x stop solution, samples were extracted with acid phenol:chloroform (5:1, (pH 4.5; Thermo Fisher Scientific), RNA products were recovered by ethanol precipitation and resuspended in NudC reaction buffer.

Full-length RNA products were incubated at 37°C for 30 min with 400 nM NudC alone (Figures 1C-D and S1B-C), 0.25 U FastAP Thermosensitive Alkaline Phosphatase alone (Figure S1B-C), or both NudC and FastAP (Figure S1B-C). Reactions were stopped by addition of an equal volume of 2x RNA loading dye (95% deionized formamide, 18 mM EDTA, 0.25% SDS, xylene cyanol, bromophenol blue, amaranth). Samples were analyzed by electrophoresis on 7.5 M urea, 1x TBE, 20% polyacrylamide gels (UreaGel System; National Diagnostics), followed by storage-phosphor imaging (Typhoon 9400 variable-mode imager; GE Life Science).

### Determination of efficiency of NCIN-mediated initiation vs. ATP-mediated initiation, (k_cat_/K_M_)_NCIN_ / (k_cat_/K_M_)_ATP_, *in vitro*: full-length product assays

For experiments in Figures 2A-B and S2, 1 *µ*M of template DNA, 1 *µ*M of mtRNAP, and 1 *µ*M transcription factor(s) (Mtf1 for *S. cerevisiae* mtRNAP; TFAM and TFB2M for human mtRNAP) were incubated at 25°C for 10 min in *Sce*-mtRNAP or human reaction buffer. A mixture containing 200 *µ*M ATP, 200 *µ*M UTP, 200 *µ*M nonradiolabeled GTP, and 6 mCi [α^32^P]-GTP at 3000 Ci/mmol and NCIN (0, 50, 100, 200, 400, 800, 1600, 3200, 6400 *µ*M) was added, and assays were incubated at 25°C for 30 min. Reactions were stopped by addition of an equal volume of 2x RNA loading dye.

Samples were analyzed by electrophoresis on 7.5 M urea, 1x TBE, 20% polyacrylamide gels (UreaGel System; National Diagnostics) supplemented with 0.2% 3-acrylamidophenylboronic acid (Boron Molecular), followed by storage-phosphor imaging (Typhoon 9400 variable-mode imager; GE Life Science).

Bands corresponding to uncapped (pppRNA) and NCIN-capped (NCIN-RNA) full-length products were quantified using ImageQuant software. The ratio of NCIN-RNA to total RNA [NCIN-RNA / (pppRNA + NCIN-RNA)] was plotted vs. the relative concentrations of NCIN vs. ATP ([NCIN] / [ATP]) on a semi-log plot (SigmaPlot) and non-linear regression was used to fit the data to the equation: y = (ax) / (b+x); where y is [NCIN-RNA / (pppRNA + NCIN-RNA)], x is ([NCIN] / [ATP]), and a and b are regression parameters. The resulting fit yields the value of x for which y = 0.5. The relative efficiency (k_cat_/K_M_)_NCIN_ / (k_cat_/K_M_)_ATP_ is equal to 1/x.

### Determination of efficiency of NCIN-mediated initiation vs. ATP-mediated initiation, (k_cat_/K_M_)_NCIN_ / (k_cat_/K_M_)_ATP_, *in vitro*: initial product assays (38)

For experiments in Figures 3C-D, and S4, 1 *µ*M of template DNA, 1 *µ*M of RNAP, and 1 *µ*M transcription factor(s) (Mtf1 for *S. cerevisiae* mtRNAP; TFAM and TFB2M for human mtRNAP; none for T7 RNAP) were incubated at 25°C for 10 min in *Sce-*mtRNAP, human mtRNAP reaction buffer, or T7 RNAP reaction buffer (40 mM Tris HCl pH 7.9, 6 mM MgCl_2_, 2 mM DTT, 2 mM Spermidine). A mixture containing 1 mM NCIN, ATP (0, 25, 50, 100, 200, 400, 800, 1600 mM), 20 *µ*M non-radiolabeled GTP, and 6 mCi [α^32^P]-GTP at 3000 Ci/mmol was added, and assays were incubated at 25°C for 30 min. Reactions were stopped by addition of an equal volume of 2x RNA loading dye. Samples were analyzed by electrophoresis on 7.5 M urea, 1x TBE, 20% polyacrylamide gels (UreaGel System; National Diagnostics) supplemented with 0.2% 3-acrylamidophenylboronic acid (Boron Molecular), followed by storage-phosphor imaging (Typhoon 9400 variable-mode imager; GE Life Science).

For experiments in Figures 2C and S3, 200 nM of fork-junction template and 500 nM RNAP (*S. cerevisiae* mtRNAP, human mtRNAP, *S. cerevisiae* RNAP II, T7 RNAP, or *E. coli* RNAP) were incubated at 25°C for 15 min in fork-junction assay buffer (10 mM Tris pH 8.0, 50 mM potassium glutamate, 10 mM MgCl_2_, 2 mM DTT, 50 ug/ml BSA). A mixture containing NCIN (1 mM NCIN for mtRNAPs, T7 RNAP, and *E. coli* RNAP; 4 mM for *S. cerevisiae* RNAP II), ATP (0, 25, 50, 100, 200, 400, 800, 1600 *µ*M for mtRNAPs and *S. cerevisiae* RNAP II; 0, 6.25, 12.5, 25, 50, 100, 200, 400 *µ*M for *E. coli* RNAP), 10 *µ*M non-radiolabeled CTP, and 6 mCi [α^32^P]-CTP (Perkin Elmer; 3000 Ci/mmol) was added, and assays were incubated at 25°C for 1 h. Reactions were stopped by addition of an equal volume of 2x RNA loading dye. Samples were analyzed by electrophoresis on 7.5 M urea, 1x TBE, 20% polyacrylamide gels (UreaGel System; National Diagnostics), followed by storage-phosphor imaging (Typhoon 9400 variable-mode imager; GE Life Science).

Bands corresponding to uncapped (pppApC) and NCIN-capped (NCINpC) initial RNA products were quantified using ImageQuant software. The ratio of NCINpC to total RNA (NCINpC / [pppApC + NCINpC]) was plotted vs. the relative concentrations of NCIN vs. ATP ([NCIN] / [ATP]) on a semi-log plot (SigmaPlot) and non-linear regression was used to fit the data to the equation: y = (ax) / (b+x); where y is [NCINpC / (pppApC + NCINpC)], x is ([NCIN] / [ATP]), and a and b are regression parameters. The resulting fit yields the value of x for which y = 0.5. The relative efficiency (k_cat_/K_M_)_NCIN_ / (k_cat_/K_M_)_ATP_ is equal to 1/x.

### Detection and quantitation of NAD^+^- and NADH-capped mitochondrial RNA *in vivo*: isolation of total cellular RNA from *S. cerevisiae*

For analysis of NAD^+^ and NADH capping during respiration, *S. cerevisiae* strain 246.1.1 [(64); *MAT*α *ura3 trp1 leu2 his4*; gift of Andrew Vershon, Rutgers University] was grown at 30°C in 25 ml YPEG (24 g Bacto-tryptone, 20 g Bacto-yeast extract, 30 mL ethanol, 3% glycerol per liter) in 125 ml flasks (Bellco) shaken at 220 rpm. When cell density reached an OD600 ~1.8 (approximately 24 hours) the cell suspension was centrifuged to collect cells (5 min, 10,000 g at 4°C), supernatants were removed, and cell pellets were resuspended in 0.8 mL RNA extraction buffer (0.5 mM NaOAc pH 5.5, 10 mM EDTA, 0.5% SDS).

For analysis of NAD^+^ and NADH capping during fermentation, *S. cerevisiae* strain 246.1.1 was grown at 30°C in 100 ml YPD [24 g Bacto-tryptone, 20 g Bacto-yeast extract, 2% (w/v) glucose per liter] in 125 ml flasks (Bellco) with airlocks to prevent oxygenation without shaking for 42 h. The cell suspension was centrifuged to collect cells (5 min, 10,000 g at 4°C), supernatants were removed, and cell pellets were resuspended in 0.8 mL RNA extraction buffer (0.5 mM NaOAc pH 5.5, 10 mM EDTA, 0.5% SDS).

To extract RNA, an equal volume of acid phenol:chloroform (5:1, pH 4.5; Thermo Fisher Scientific) was added to cells in resuspension buffer and mixed by vortexing for 10 s. Samples were incubated at 65°C for 5 min, -80°C for 5 min, then centrifuged (15 min, 21,000 g, 4°C) to separate the aqueous and organic phases. The aqueous phase was collected and acid phenol:chloroform extraction was performed two more times on this solution. RNA transcripts were recovered by ethanol precipitation and resuspended in RNase free H_2_O.

### Detection and quantitation of NAD^+^- and NADH-capped mitochondrial RNA *in vivo*: isolation of total cellular RNA from human cells

Human embryonic kidney HEK293T cells (obtained from ATCC) were maintained under 5% CO_2_ at 37°C in DMEM medium (Thermo Fisher Scientific) supplemented with 10% fetal bovine serum (Atlanta Biologicals), 100 units/ml penicillin, and 100 *µ*g/ml streptomycin. HEK293T cells were seeded in 100 mm tissue-culture treated plates and grown for 72 h at 37°C or seeded in 100 mm tissue-culture treated plates, grown for 24 h at 37°C, treated with 5 nM FK866 (Sigma Aldrich), and grown for an additional 48 h at 37°C. Total cellular RNA was isolated with TRIzol Reagent according to the manufacture’s protocol (Thermo Fisher Scientific).

### Detection and quantitation of NAD^+^- and NADH-capped mitochondrial RNA *in vivo*: DNAzyme cleavage

For analysis of NCIN capping of *S. cerevisiae* mitochondrial RNA, 40 *µ*g of total cellular RNA was mixed with 1 *µ*M DNAzyme (JB557 for *S. cerevisiae* COX2 RNA; JB526 for *S. cerevisiae* 21S RNA) in buffer containing 10 mM Tris pH 8.0, 50 mM NaCl, 2 mM DTT, and 10 mM MgCl_2_ (total volume 50 *µ*l). When present, NudC was added to 400 nM. Reactions were incubated for 60 min at 37°C and 100 *µ*l of 1.5x stop solution and 500 *µ*l ethanol was added. Samples were centrifuged (30 min, 21,000 g, 4°C), the supernatant removed, and the pellet resuspended in 2x RNA loading dye.

For analysis of NCIN capping of human mitochondrial RNA, 40 *µ*g of total cellular RNA was mixed with 1 *µ*M DNAzyme (JB559 for human LSP-generated RNA) in buffer containing 10 mM Tris pH 8.0, 50 mM NaCl, 2 mM DTT (total volume 50 *µ*l). Samples were heated to 85°C for 5 minutes, cooled to 37°C. MgCl_2_ was added to a final concentration of 10 mM and, when present, NudC was added to 400 nM. Reactions were incubated for 60 min at 37°C and 100 *µ*l of 1.5x stop solution and 500 *µ*l ethanol was added. Samples were centrifuged (30 min, 21,000 g, 4°C), the supernatant removed, and the pellet resuspended in 2x RNA loading dye.

To prepare synthetic RNA standards non-radiolabeled full-length RNA products were generated by *in vitro* transcription reactions containing 1 *µ*M of template DNA, 1 *µ*M of mtRNAP, 1 *µ*M transcription factor(s), 1 mM initiating nucleotide (ATP, NAD^+^ or NADH), 200 *µ*M GTP, 200 *µ*M UTP, 200 *µ*M CTP, and 20 *µ*M ATP. Reactions were incubated for 60 min at 25°C, stopped by addition of 1.5x stop solution, extracted with acid phenol:chloroform (5:1, pH 4.5; Thermo Fisher Scientific) and ethanol precipitated. Full-length RNAs were resuspended in buffer containing 10 mM Tris pH 8.0, 50 mM NaCl, 2 mM DTT, and 10 mM MgCl_2_ and treated with DNAzyme as described above.

### Detection and quantitation of NAD^+^- and NADH-capped mitochondrial RNA *in vivo*: hybridization with a radiolabeled oligodeoxyribonucleotide probe

NCIN capping of DNAzyme-generated subfragments of mitochondrial RNA were analyzed by a procedure consisting of: (i) electrophoresis on 7.5 M urea, 1x TBE, 10% polyacrylamide gels supplemented with 0.2% 3-acrylamidophenylboronic acid (Boron Molecular); (ii) transfer of nucleic acids to a Nytran supercharge nylon membrane (GE Healthcare Life Sciences) using a semidry transfer apparatus (Bio-Rad); (iii) immobilization of nucleic acids by UV crosslinking; (iv) incubation with a ^32^P-labelled oligodeoxyribonucleotide probe complementary to target RNAs (JB555, COX2 RNA; JB525, 21S RNA; JB515, LSP-derived RNA; ^32^P-labelled using T4 polynucleotide kinase and [γ^32^P]-ATP [Perkin Elmer]); (v) high-stringency washing, procedures as in (65); and (vi) storage-phosphor imaging (Typhoon 9400 variable-mode imager; GE Life Science).

Bands corresponding to uncapped and NCIN-capped DNAzyme-generated subfragments were quantified using ImageQuant software. The percentages of uncapped RNA (5’-ppp), NAD^+^-capped RNA (5’-NAD^+^), or NADH-capped RNA (5’-NADH) to total RNA were determined from three biological replicates.

### Determination of NAD(H) levels in human HEK293T cells

Cells were lysed in 400 μl of NAD/H Extraction Buffer (NAD/H Quantitation Kit, Sigma-Aldrich) and total proteins were extracted with two freeze-thaw cycles (20 min on dry ice, 10 min at RT). Lysates were centrifuged (10 min, 13,000 g, 4°C), the supernatant was isolated, and the protein concentration was determined. NAD(H) was measured using a NAD/H Quantitation Kit (Sigma) from a volume of supernatant containing 10 μg protein.

Molecules of NAD(H) per cell were calculated using a value of 300 pg total protein per cell. The cellular concentration of NAD(H) was then estimated by using a cellular volume of ~ 1200 *µ*m^3^ for HEK293T cells and assuming a homogenous distribution of NAD(H) in the cell. The volume of HEK293T cells was calculated using a value of 13 *µ*m for the diameter of HEK293T cells, assuming trypsinized cells adopt a spherical shape.

## References

1. Hofer K – Jaschke A (2018) Epitranscriptomics: RNA Modifications in Bacteria and Archaea. Microbiol Spectr 6(3).

2. Shuman S (2015) RNA capping: progress and prospects. RNA 21(4):735-737.

3. Ramanathan A, Robb GB, – Chan SH (2016) mRNA capping: biological functions and applications. Nucleic Acids Res 44(16):7511-7526.

4. Jaschke A, Hofer K, Nubel G, – Frindert J (2016) Cap-like structures in bacterial RNA and epitranscriptomic modification. Current opinion in microbiology 30:44-49.

5. Furuichi Y – Shatkin AJ (2000) Viral and cellular mRNA capping: past and prospects. Adv Virus Res 55:135-184.

6. Shuman S (1995) Capping enzyme in eukaryotic mRNA synthesis. Prog Nucleic Acid Res Mol Biol 50:101-129.

7. Shatkin AJ (1976) Capping of eucaryotic mRNAs. Cell 9(4 PT 2):645-653.

8. Wei CM, Gershowitz A, – Moss B (1975) Methylated nucleotides block 5’ terminus of HeLa cell messenger RNA. Cell 4(4):379-386.

9. Chen YG, Kowtoniuk WE, Agarwal I, Shen Y, – Liu DR (2009) LC/MS analysis of cellular RNA reveals NAD-linked RNA. Nature chemical biology 5(12):879-881.

10. Walters RW, et al. (2017) Identification of NAD+ capped mRNAs in Saccharomyces cerevisiae. Proc Natl Acad Sci U S A 114(3):480-485.

11. Jiao X, et al. (2017) 5’ End Nicotinamide Adenine Dinucleotide Cap in Human Cells Promotes RNA Decay through DXO-Mediated deNADding. Cell 168(6):1015-1027 e1010.

12. Cahova H, Winz ML, Hofer K, Nubel G, – Jaschke A (2015) NAD captureSeq indicates NAD as a bacterial cap for a subset of regulatory RNAs. Nature 519(7543):374-377.

13. Frindert J, et al. (2018) Identification, Biosynthesis, and Decapping of NAD-Capped RNAs in B. subtilis. Cell Rep 24(7):1890-1901 e1898.

14. Ghosh A – Lima CD (2010) Enzymology of RNA cap synthesis. Wiley Interdiscip Rev RNA 1(1):152-172.

15. Martinez-Rucobo FW, et al. (2015) Molecular Basis of Transcription-Coupled Pre-mRNA Capping. Mol Cell 58(6):1079-1089.

16. Shuman S (2001) Structure, mechanism, and evolution of the mRNA capping apparatus. Prog Nucleic Acid Res Mol Biol 66:1-40.

17. Bird JG, et al. (2016) The mechanism of RNA 5’ capping with NAD+, NADH and desphospho-CoA. Nature 535(7612):444-447.

18. Julius C, Riaz-Bradley A, – Yuzenkova Y (2018) RNA capping by mitochondrial and multi-subunit RNA polymerases. Transcription.

19. Barvik I, Rejman D, Panova N, Sanderova H, – Krasny L (2017) Non-canonical transcription initiation: the expanding universe of transcription initiating substrates. FEMS Microbiol Rev 41(2):131-138.

20. Julius C – Yuzenkova Y (2017) Bacterial RNA polymerase caps RNA with various cofactors and cell wall precursors. Nucleic Acids Res 45(14):8282-8290.

21. Vvedenskaya IO, et al. (2018) CapZyme-Seq Comprehensively Defines Promoter-Sequence Determinants for RNA 5’ Capping with NAD^+^. Mol Cell 70(3):553-564 e559.

22. Winz ML, et al. (2017) Capture and sequencing of NAD-capped RNA sequences with NAD captureSeq. Nature protocols 12(1):122-149.

23. Cramer P (2002) Multisubunit RNA polymerases. Curr Opin Struct Biol 12(1):89-97.

24. Darst SA (2001) Bacterial RNA polymerase. Curr Opin Struct Biol 11(2):155-162.

25. Ebright RH (2000) RNA polymerase: structural similarities between bacterial RNA polymerase and eukaryotic RNA polymerase II. J Mol Biol 304(5):687-698.

26. Werner F – Grohmann D (2011) Evolution of multisubunit RNA polymerases in the three domains of life. Nat Rev Microbiol 9(2):85-98.

27. Hillen HS, Temiakov D, – Cramer P (2018) Structural basis of mitochondrial transcription. Nat Struct Mol Biol 25(9):754-765.

28. Ringel R, et al. (2011) Structure of human mitochondrial RNA polymerase. Nature 478(7368):269-273.

29. Cermakian N, Ikeda TM, Cedergren R, – Gray MW (1996) Sequences homologous to yeast mitochondrial and bacteriophage T3 and T7 RNA polymerases are widespread throughout the eukaryotic lineage. Nucleic Acids Res 24(4):648-654.

30. Masters BS, Stohl LL, – Clayton DA (1987) Yeast mitochondrial RNA polymerase is homologous to those encoded by bacteriophages T3 and T7. Cell 51(1):89-99.

31. Cheetham GM – Steitz TA (2000) Insights into transcription: structure and function of single-subunit DNA-dependent RNA polymerases. Curr Opin Struct Biol 10(1):117-123.

32. Sousa R (1996) Structural and mechanistic relationships between nucleic acid polymerases. Trends in biochemical sciences 21(5):186-190.

33. McAllister WT – Raskin CA (1993) The phage RNA polymerases are related to DNA polymerases and reverse transcriptases. Mol Microbiol 10(1):1-6.

34. Deshpande AP – Patel SS (2014) Interactions of the yeast mitochondrial RNA polymerase with the +1 and +2 promoter bases dictate transcription initiation efficiency. Nucleic Acids Res 42(18):11721-11732.

35. Sologub M, Litonin D, Anikin M, Mustaev A, – Temiakov D (2009) TFB2 is a transient component of the catalytic site of the human mitochondrial RNA polymerase. Cell 139(5):934-944.

36. Deana A, Celesnik H, – Belasco JG (2008) The bacterial enzyme RppH triggers messenger RNA degradation by 5’ pyrophosphate removal. Nature 451(7176):355-358.

37. Hofer K, et al. (2016) Structure and function of the bacterial decapping enzyme NudC. Nature chemical biology 12(9):730-734.

38. Bird JG, Nickels BE, – Ebright RH (2017) RNA Capping by Transcription Initiation with Non-canonical Initiating Nucleotides (NCINs): Determination of Relative Efficiencies of Transcription Initiation with NCINs and NTPs. Bio-protocol 7(12).

39. Igloi GL – Kossel H (1987) Use of boronate-containing gels for electrophoretic analysis of both ends of RNA molecules. Methods Enzymol 155:433-448.

40. Igloi GL – Kossel H (1985) Affinity electrophoresis for monitoring terminal phosphorylation and the presence of queuosine in RNA. Application of polyacrylamide containing a covalently bound boronic acid. Nucleic Acids Res 13(19):6881-6898.

41. Nubel G, Sorgenfrei FA, – Jaschke A (2017) Boronate affinity electrophoresis for the purification and analysis of cofactor-modified RNAs. Methods 117:14-20.

42. Joyce GF (2001) RNA cleavage by the 10-23 DNA enzyme. Methods Enzymol 341:503-517.

43. Canelas AB, van Gulik WM, – Heijnen JJ (2008) Determination of the cytosolic free NAD/NADH ratio in Saccharomyces cerevisiae under steady-state and highly dynamic conditions. Biotechnol Bioeng 100(4):734-743.

44. Bekers KM, Heijnen JJ, – van Gulik WM (2015) Determination of the in vivo NAD:NADH ratio in *Saccharomyces cerevisiae* under anaerobic conditions, using alcohol dehydrogenase as sensor reaction. Yeast 32(8):541-557.

45. Hasmann M – Schemainda I (2003) FK866, a highly specific noncompetitive inhibitor of nicotinamide phosphoribosyltransferase, represents a novel mechanism for induction of tumor cell apoptosis. Cancer Res 63(21):7436-7442.

46. Khan JA, Tao X, – Tong L (2006) Molecular basis for the inhibition of human NMPRTase, a novel target for anticancer agents. Nat Struct Mol Biol 13(7):582-588.

47. Chen WW, Freinkman E, Wang T, Birsoy K, – Sabatini DM (2016) Absolute Quantification of Matrix Metabolites Reveals the Dynamics of Mitochondrial Metabolism. Cell 166(5):1324-1337 e1311.

48. Cambronne XA, et al. (2016) Biosensor reveals multiple sources for mitochondrial NAD(+). Science 352(6292):1474-1477.

49. Biswas TK (1999) Nucleotide sequences surrounding the nonanucleotide promoter motif influence the activity of yeast mitochondrial promoter. Biochemistry 38(30):9693-9703.

50. Taanman JW (1999) The mitochondrial genome: structure, transcription, translation and replication. Biochim Biophys Acta 1410(2):103-123.

51. Chang DD – Clayton DA (1984) Precise identification of individual promoters for transcription of each strand of human mitochondrial DNA. Cell 36(3):635-643.

52. Thomason MK, et al. (2015) Global transcriptional start site mapping using differential RNA sequencing reveals novel antisense RNAs in Escherichia coli. Journal of bacteriology 197(1):18-28.

53. Haberle V, et al. (2014) Two independent transcription initiation codes overlap on vertebrate core promoters. Nature 507(7492):381-385.

54. Saito TL, et al. (2013) The transcription start site landscape of *C. elegans*. Genome Res 23(8):1348-1361.

55. Chen RA, et al. (2013) The landscape of RNA polymerase II transcription initiation in C. elegans reveals promoter and enhancer architectures. Genome Res 23(8):1339-1347.

56. Tsuchihara K, et al. (2009) Massive transcriptional start site analysis of human genes in hypoxia cells. Nucleic Acids Res 37(7):2249-2263.

57. Tang GQ, Paratkar S, – Patel SS (2009) Fluorescence mapping of the open complex of yeast mitochondrial RNA polymerase. The Journal of biological chemistry 284(9):5514-5522.

58. Paratkar S – Patel SS (2010) Mitochondrial transcription factor Mtf1 traps the unwound non-template strand to facilitate open complex formation. The Journal of biological chemistry 285(6):3949-3956.

59. Ramachandran A, Basu U, Sultana S, Nandakumar D, – Patel SS (2017) Human mitochondrial transcription factors TFAM and TFB2M work synergistically in promoter melting during transcription initiation. Nucleic Acids Res 45(2):861-874.

60. Yakubovskaya E, et al. (2014) Organization of the human mitochondrial transcription initiation complex. Nucleic Acids Res 42(6):4100-4112.

61. Jia Y, Kumar A, – Patel SS (1996) Equilibrium and stopped-flow kinetic studies of interaction between T7 RNA polymerase and its promoters measured by protein and 2-aminopurine fluorescence changes. The Journal of biological chemistry 271(48):30451-30458.

62. Artsimovitch I, Svetlov V, Murakami KS, – Landick R (2003) Co-overexpression of *Escherichia coli* RNA polymerase subunits allows isolation and analysis of mutant enzymes lacking lineage-specific sequence insertions. The Journal of biological chemistry 278(14):12344-12355.

63. Barnes CO, et al. (2015) Crystal Structure of a Transcribing RNA Polymerase II Complex Reveals a Complete Transcription Bubble. Mol Cell 59(2):258-269.

64. Tatchell K, Nasmyth KA, Hall BD, Astell C, – Smith M (1981) In vitro mutation analysis of the mating-type locus in yeast. Cell 27(1 Pt 2):25-35.

65. Goldman SR, Nair NU, Wells CD, Nickels BE, – Hochschild A (2015) The primary sigma factor in *Escherichia coli* can access the transcription elongation complex from solution in vivo. Elife 4.

